# Disentangling Mixed Classes of Covariability in Large-Scale Neural Data

**DOI:** 10.1101/2023.03.01.530616

**Authors:** Arthur Pellegrino, Heike Stein, N Alex Cayco-Gajic

## Abstract

Recent work has argued that large-scale neural recordings are often well described by low-dimensional ‘latent’ dynamics identified using dimensionality reduction. However, the view that task-relevant variability is shared across neurons misses other types of structure underlying behavior, including stereotyped neural sequences or slowly evolving latent spaces. To address this, we introduce a new framework that simultaneously accounts for variability that is shared across neurons, trials, or time. To identify and demix these covariability classes, we develop a new unsupervised dimensionality reduction method for neural data tensors called sliceTCA. In three example datasets, including motor cortical dynamics during a classic reaching task and recent multi-region recordings from the International Brain Laboratory, we show that sliceTCA can capture more task-relevant structure in neural data using fewer components than traditional methods. Overall, our theoretical framework extends the classic view of low-dimensional population activity by incorporating additional classes of latent variables capturing higher-dimensional structure.

## 1 Introduction

Neural activity varies in relation to fluctuations in the environment, slow changes in circuitry, and heterogeneous cell properties, creating variability across neurons, time, and trials. Recent work has emphasized that trial-to-trial variability is often correlated across large populations of neurons [Cunningham and Yu, 2014], generating low-dimensional representations of sensory or behavioral variables. Indeed, analyzing shared variability across neurons has led to key insights into the information encoded and computations performed by neural circuits [Panzeri et al., 2022, Jazayeri and Ostojic, 2021]. Such findings have driven an increase in the popularity of dimensionality reduction methods, such as principal component analysis (PCA), which seek to capture structure in neural data by identifying covarying population-wide patterns. More recent work has advocated instead for applying tensor-based methods, such as tensor component analysis (TCA), that distinguish between changes in neural dynamics that occur over fast (within-trial) and slow (between-trial) timescales [Williams et al., 2018, Harshman et al., 1970, Carroll and Chang, 1970]. In both of these approaches, neural activity is assumed to be constrained to a low-dimensional neural subspace (defined by a set of latent variables) that is fixed over the course of an experiment.

However, this picture of latent variables fails to account for some forms of shared variability in neural circuits. First, not all population dynamics are described by a fixed covariance structure. For example, many brain areas produce temporal sequences in which the latency of activation varies from neuron to neuron, but which are highly stereotyped across conditions [Seely et al., 2016, Pastalkova et al., 2008, Peters et al., 2014, Okubo et al., 2015, Harvey et al., 2012]. Second, the neural encoding weights for a given sensory stimulus may change over trials due to adaptation, learning, [Hennig et al., 2021], or representational drift [Rule et al., 2019, Driscoll et al., 2017, Schoonover et al., 2021]. Because methods such as PCA or TCA look for covariability across neurons, they may miss additional forms of variability that are instead shared across time or across trials.

To address this, we introduce slice tensor component analysis (sliceTCA), an unsupervised dimensionality reduction method that is able to identify and disentangle latent variables belonging to three different classes of covariability (defined as variability shared across neurons, time, or trials) that are mixed within the same dataset. This property contrasts sliceTCA from matrix factorizations like PCA which capture a single covariability class at a time, and from TCA which identifies components constrained to all of them simultaneously. As a result, we show that sliceTCA can capture more structure in fewer components than either of these methods. Based on theoretical and practical considerations of the sliceTCA decomposition, we develop an analysis pipeline for model selection, optimization, and visualization that is implemented in a readily applicable Python library.

After validating our method on simulated data, we illustrate the utility of sliceTCA in three large-scale neural datasets. First, we demonstrate that different covariability classes encode distinct behaviorally relevant neural dynamics in motor cortical recordings in non-human primates [Churchland et al., 2012]. Next, in simultaneous imaging data from cortical and cerebellar populations during a cued motor task [Wagner et al., 2019], we show that sliceTCA untangles task-relevant manifolds by taking into account covariability across trials. Finally, we analyze a recent dataset from the International Brain Laboratory [IBL et al., 2022] and show that sliceTCA disentangles region-specific covariability classes across visual cortex, hippocampus, thalamus, and the midbrain. We then provide a geometric intuition for how neural population activity is shaped by latent variables belonging to the three different covariability classes. Together, these results demonstrate the necessity of extending the traditional view of latent variables and neural covariability to uncover higher-dimensional latent structure. With sliceTCA, we propose a novel, unsupervised dimensionality reduction method that uncovers co-existing classes of behaviorally relevant covariability in neural datasets.

## 2 Results

### 2.1 Overview of sliceTCA

SliceTCA is an unsupervised dimensionality reduction method that generalizes matrix factorizations, a class of methods which includes PCA and non-negative matrix factorization (NMF). Matrix factorizations approximate a data matrix **X** as a sum of *R* components:

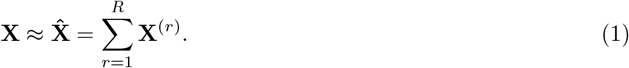

In neuroscience, **X** is generally a matrix of size *N* × *KT* containing the activity of *N* neurons recorded over *K* trials, each containing *T* timepoints. Each component **X**^(*r*)^ is a rank-1 matrix defined by a set of *R* neural factors describing different activation patterns across the population, and a set of *R* corresponding temporal factors describing how the strength of these patterns evolves over the course of experiment (Figure 1a). By constraining each component to a rank-1 matrix, these methods capture shared variability across neurons.

**Figure 1:**
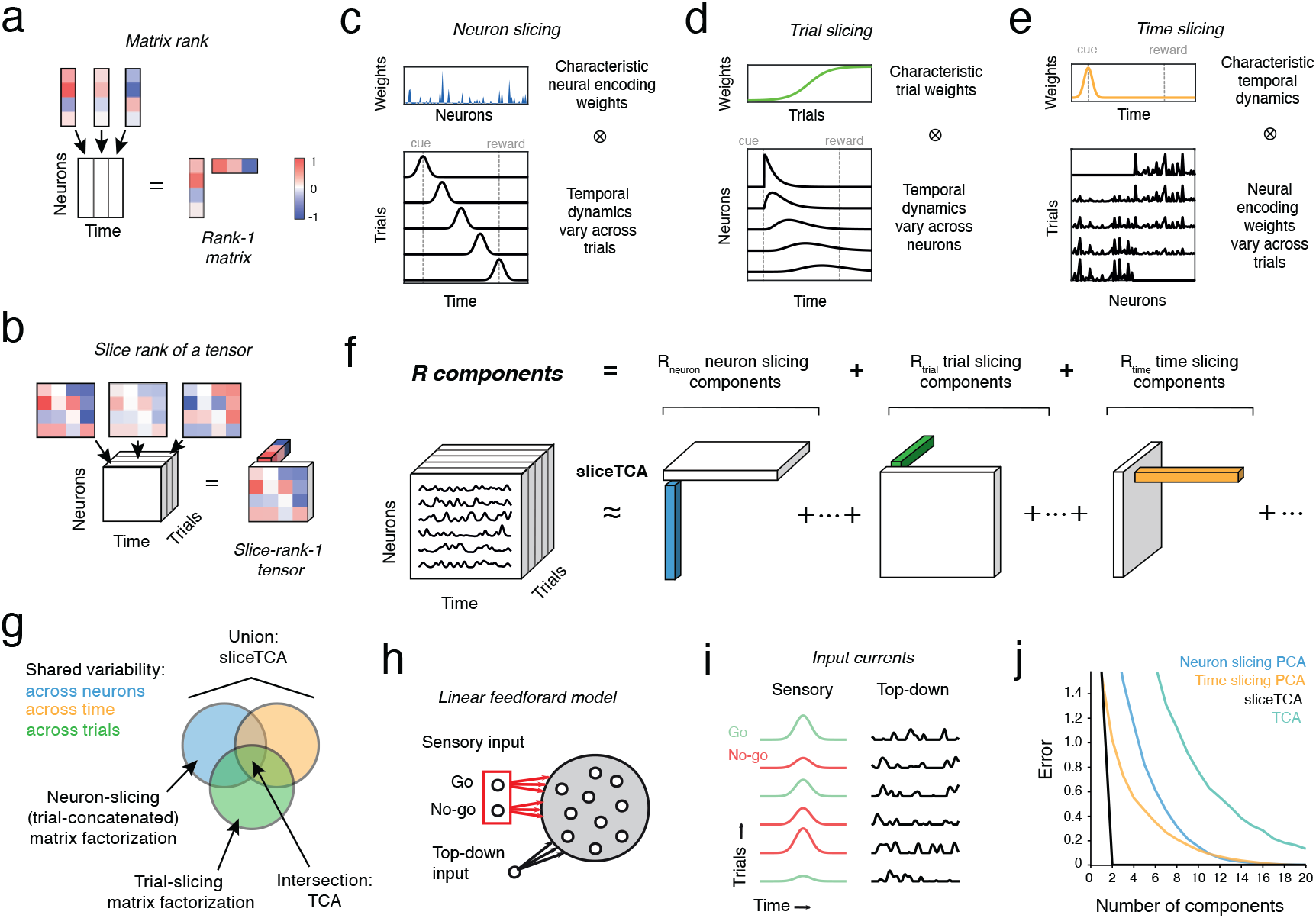
SliceTCA demixes shared variability across neurons, time, and trials. **a.** Schematic representation of a rank 1 matrix. Each column of the matrix is a scaled version of the same vector. Equivalently, the matrix can be written as the outer product of that same column vector and a row vector representing the scaling weights. **b.** Schematic representation of a slice-rank-1 tensor. Each “slice” of the tensor is a scaled version of the same matrix. The tensor can be written as an outer product of that matrix (a “slice”) and a vector representing the scaling weights. **c.** Example of a slice-rank-1 tensor that is an outer product of a neuron loading vector and a time-trial slice. This component represents a latent variable with a fixed neural encoding but whose temporal profile changes from trial to trial. **d.** Example slice-rank-1 tensor that is an outer product of a trial loading vector and a neuron-time slice, representing a latent variable that scales in amplitude over trials but which has neuron specific dynamics within a trial. **e.** Example slice-rank-1 tensor that is an outer product of a time loading vector and a neuron-trial slice, representing a latent variable with a characteristic temporal profile within a trial, but whose neural encoding weights change over trials. **f.** SliceTCA approximates the data tensor as a low-slice-rank approximation. Each component is described as a slice-rank-1 tensor, which can be one of three types: neuron-, trial-, or time-slicing, corresponding to the examples in (c-e). **g.** Schematic of the three covariability classes captured by sliceTCA. Matrix factorization methods like PCA only capture a single covariability class at a time, depending on how the data tensor is unfolded into matrix form. TCA requires the variability captured by each component to be shared across neurons, trials, and time simultaneously (i.e., in the intersection of the three classes). In contrast, sliceTCA represents the union of these three classes. **h.** Schematic of a toy model of perceptual learning during a Go/No-go task. On each trial, a population of linear neurons receives two inputs: (1) sensory input from one of two upstream sources representing the presentation of the Go or No-go stimulus, and (2) top-down modulation representing stimulus-independent factors. Red indicates plastic synapses (Go/No-go weights potentiate/depress over Go/No-go trials, respectively). **i.** Evolution of inputs over trials. Go/No-go inputs increase/decrease in strength over trials, while top-down inputs vary from trial to trial. Since the neurons are linear, their activities will be a linear combination of these two input sources. **j.** Loss (mean squared error) curves as a function of the number of components for different methods.

However, this approach to neural dimensionality reduction has two limitations. First, by concatenating trials together to structure the data into a matrix, they do not distinguish between rapid dynamics within a trial and slower changes across trials [Williams et al., 2018]. Second, not all population activity is well described by shared variability across neurons. For example, motor cortical dynamics may be better described by stereotyped sequences of neural activation which are shared across trials of the same condition [Seely et al., 2016]. These limitations can be addressed by structuring the data into an *N* × *T* × *K tensor* which can be similarly decomposed following equation 1 into a low-rank *tensor* approximation. For this, we must generalize the concept of a rank-1 matrix to tensors. Different definitions of the tensor rank will capture different forms of structure in the data.

Here, we present *sliceTCA,* a novel tensor decomposition that is based on the *slice rank* [Tao and Sawin, 2016] (Methods). A rank-1 matrix is defined as the outer product of two vectors, so that each column of the matrix is a scaled version of the same column vector (Figure 1a). Similarly, a slice-rank-1 tensor is defined as the outer product of a vector and a matrix (or “slice”; Figure 1b). Depending on how the tensor is sliced, these components can capture three different classes of covariability (across neurons, trials, or time).

To gain intuition on this, we may consider each slice type separately. First, a neuron-slicing component is described by a vector of characteristic neural weights and a matrix describing how the temporal dynamics for that component changes across time and trials (Figure 1c). This component therefore captures variability which is shared across neurons, but which is unconstrained across time and trials. This is exactly the same class of covariability that is captured by common applications of matrix factorization methods in which the data tensor is reshaped or ‘unfolded’ into a *N* × *KT* matrix (sometimes referred to as ‘trial-concatenated’ matrix factorization, Supplementary Figure 1a). In contrast, the other two slice types lead to different assumptions about the source of covariability in the data. The trial-slicing components instead capture shared variability across trials, e.g., stereotyped neuron-specific temporal dynamics that vary together in amplitude over trials (Figure 1d). Finally, the time-slicing components identify shared variability over time. This could represent common dynamics whose neural encoding weights change from trial to trial, e.g., due to learning, adaptation, or drift (Figure 1e).

If only one of these three slice types were fitted, sliceTCA would be equivalent to a matrix factorization on the respective unfolding of the data tensor (Supplementary Figure 1a-c). Indeed, previous work has argued for performing PCA on different unfoldings of the data tensor to identify the slice type that gives the best approximation [Seely et al., 2016]. Crucially, sliceTCA differs from this approach by fitting all three slice types *simultaneously,* thereby demixing different covariability classes that may be combined within the same dataset (Figure 1f,g). SliceTCA is also related to, yet distinct from TCA (also known as CANDECOMP/PARAFAC or CP decomposition) [Williams et al., 2018, Harshman et al., 1970, Carroll and Chang, 1970]. TCA constrains each component to be described by the outer product of three vectors of neural, trial, and temporal factors, which requires that each component must lie in the intersection of all three covariability classes (Figure 1g; Methods). As a result, sliceTCA is able to capture more structure in the data with fewer components as compared to other methods.

### 2.2 Mixed variability in a simple model of perceptual learning

Before applying sliceTCA to data, we first illustrate how mixed covariability classes could emerge in neural circuits. We built a toy model of sensory cortex during a Go/No-go task (Figure 1h). In this model, a population of linear cortical neurons received two sources of input in the context of a Go/No-go task (Figure 1i; Supplementary Figure 2a). First, all neurons received sensory input representing the presented stimulus (either Go or No-go). The projection weights were plastic and subject to potentiation or depression (for the Go / No-go stimuli, respectively), in line with evidence of enhanced sensitivity to target stimuli in sensory cortex during perceptual learning [Poort et al., 2015]. The evolution of neural weights over trials was stochastic with heterogeneous learning rates, creating variability across neurons (Supplementary Figure 2b; Methods). Second, all neurons also received input representing top-down modulatory processes such as arousal or behavior which may vary from trial to trial, but which were not directly related to the task [Vinck et al., 2015]. In this linear model, each neuron’s activity is simply the summation of its sensory and top-down input currents (Figure 1i).

From these minimal assumptions, the two input sources represent two different classes of covariability. First, the sensory input has a characteristic temporal dynamics that is time-locked to stimulus presentation with neural encoding weights that vary over trials due to heterogeneity in the learning dynamics. In contrast, the top-down input source is characterized by fixed, non-plastic neural encoding weights but with trial-to-trial variability in the temporal dynamics. The resulting population activity has slice rank of two, as the sum of one time-slicing component (sensory input) and one neuron-slicing component (top-down input). Indeed, sliceTCA is able to capture the two ground truth components (Figure 1j, Supplementary Figure 2c). On the other hand, PCA and TCA require significantly more components to capture the variability (Figure 1j). SliceTCA outperformed PCA and TCA even when white noise was added to the data (Supplementary Figure 2d,e). Together these results show that mixed covariability classes can emerge from minimal assumptions about heterogeneity in neural circuits, and that they can be disentangled using sliceTCA.

### 2.3 Task relevant information is distributed across slice types in motor cortical reaching dynamics

Based on the results of our toy model, we predicted that different slice types could capture different kinds of behaviorally relevant dynamics in neural data. We tested this hypothesis in a dataset comprising primary motor cortical (M1) and premotor cortical (PMd) populations recorded simultaneously during maze reaching and classic center-out (no maze) reaching tasks (Figure 2a, hand position). To quantify decoding performance, we linearly mapped population activity onto hand velocity (Methods). As a benchmark, we first mapped trial-averaged raw neural data on kinematic trajectories, revealing a close match between behavior and neural activity (Figure 2a, trial-averaged raw data). However, when we attempted to decode hand trajectories based on individual trials, we observed significant trial-to-trial variability that corresponded poorly to kinematic data (Figure 2a, raw data).

**Figure 2:**
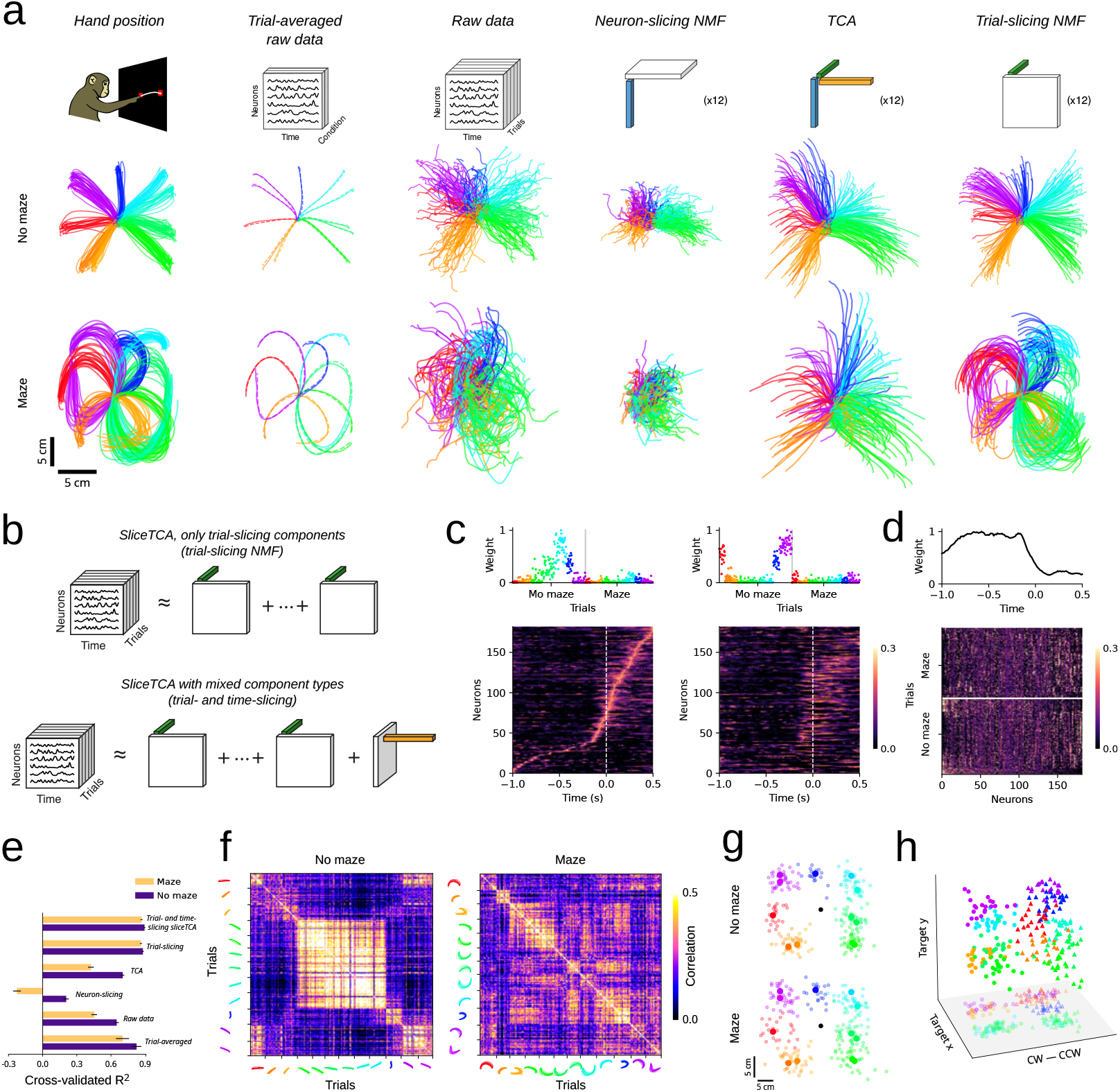
Time and trial slicing components identify preparatory and kinematic information in motor cortical dynamics, respectively. **a.** Behavioral and motor cortical dynamics (n = 182 neurons from M1/PMd) during a classic center-out reaching task with straight reaches (top) and curved maze reaches (bottom, modified from Churchland et al. [2012]). Different colours indicate different reach directions. *Hand position.* Hand positions during the experiment. *Trial-averaged raw data*. Condition-wise trial-averaged reaches (dashed lines) vs. condition-averaged neural population activity (solid lines), projected onto the 2D subspace that best matches hand trajectories. *Raw data.* Trial-by-trial mapping of raw population activity onto hand trajectories *Neuron-slicing.* Trial-by-trial mapping of denoised population activity onto hand trajectories (neuron-slicing NMF, 12 components; equivalent to NMF performed on the trial-concatenated data matrix). *TCA.* Trial-by-trial mapping of denoised population activity (TCA, 12 components). *Trial-slicing*. Trial-by-trial mapping of denoised population activity (trial-slicing NMF, 12 components) onto hand trajectories. **b.** Schematics of sliceTCA models with multiple components of the same slice type vs. a model with mixed slice types. **c.** Two example trial-slicing components, ordered by peak activation times of the first component. Sequential patterns distinguished specific reach conditions (here, upper left vs. lower right) **d.** The single time-slicing component has high temporal weight preceding movement onset, as well as condition-specific neural weights in the slice. **e.** *R*^2^ of 5-fold cross-validated velocity decoding in each model. **f.** In PMd, correlations between neural weights on the time-slicing component were high for pairs of trials with similar reach direction and curvature, and low for dissimilar reaches. **g.** Mapping of average activity in the time-slicing component before movement onset (0.75 — 0s pre-onset) onto reach targets reveals a strong association (*R*^2^ = 0.95 and *R*^2^ = 0.91, center-out vs. curved reaches) **h.** Partially reconstructed activity from the time-slicing component, projected into a three-dimensional subspace identified to maximally separate clockwise vs. counter-clockwise movements and target x and y positions. Data points are clustered according to both reach direction and curvature, indicating that the time-slicing component encodes information about the dynamics of the upcoming movement (dots indicate clockwise, triangles counter-clockwise reaches).

We reasoned that single-trial kinematic information might be present in the data, but obscured by behaviorally irrelevant neural variability. If true, then the decoder should perform significantly better on properly denoised data. To test this, we first used a common approach of fitting a low-rank approximation using non-negative matrix factorization (NMF, *R* =12 components) to the *N* × (*TK*) matrix of trial-concatenated neural activity (‘neuron-unfolded’ data). Surprisingly, this actually decreased the performance of the decoder (Figure 2a, neuron-slicing NMF), suggesting that the variability discarded by this denoising procedure contains information about hand kinematics. We wondered whether better performance could be obtained with a method that explicitly identifies shared variability across trials. Indeed, TCA-denoised data displayed a better match to the hand kinematics (*R* = 12 components, Figure 2a, TCA). Yet, by constraining the decomposition to be low tensor rank and thus also discarding temporal variability across neurons, TCA is not able to reconstruct neural sequences at a sufficiently high temporal resolution to allow for precise behavioral readout.

By performing TCA and NMF on the neuron-unfolded data tensor, we have assumed that behaviourally-relevant variability in the data is shared across neurons (Figure 1g). However, previous work has emphasized that dynamics in motor regions are better described by stereotyped sequences of activation that are shared across conditions [Seely et al., 2016, Mackevicius et al., 2019]. Following this intuition, we performed the same decoding analysis on denoised trial-unfolded data, where a *T* × (*NK*) matrix is approximated using NMF (*R* = 12 components). Remarkably, this simple change in denoising strategy resulted in a significantly better match between trial-to-trial variability in the data and in the hand kinematics (Figure 2a, trial-slicing NMF). We further validated that the components obtained by trial-slicing NMF corresponded to reach-tuned sequences whose temporal orderings were reproducible across held-out data (Supplementary Figure 3). These results reveal that in this dataset, behaviorally relevant information was encoded in neural sequences shared over trials, rather than by shared variability across neurons.

Trial- and neuron-concatenated NMF constitute two special cases of non-negative sliceTCA, where either neuron-slicing components or trial-slicing components exclusively are fitted. Therefore, we next asked whether we could identify additional information in the data by demixing different classes of covariability with sliceTCA. Previous work has identified preparatory signals in the premotor cortex that indicate the dynamics of the upcoming movement [Shenoy et al., 2013]. We therefore hypothesized that we could capture preparatory signals in a time-slicing component with shared ramping dynamics, and neural encoding weights encoding reach targets and curvature on a trial-by-trial basis.

Towards this end, we used sliceTCA to add a single time-slicing component to the previous model with 12 trial-slicing components (Figure 2b; Supplementary Figure 4). In both the trial-slicing NMF model and the sliceTCA model with mixed slice types, the trial-slicing components identified sequential neural dynamics for similar reach conditions which seemed to be continuously tuned to target angles (Figure 2c; Supplementary Figure 4b). Decoding from these trial-slicing components (in either the mixed or the unmixed model) led to significantly better performance as compared to the neuron-slicing and TCA models (Figure 2e; Supplementary Figure 5). The trial-slicing partial reconstruction from sliceTCA mapped slightly better onto hand kinematics in the mixed model than in the trial-slicing only model (Figure 2e, *p* < 0.001, Wilcoxon signed rank test). Intriguingly, while the single time-slicing component mapped poorly onto hand kinematics (Supplementary Figure 4a), it identified shared dynamics that peaked around 100 ms before movement onset (Figure 2d), consistent with a preparatory movement signal.

If the time-slicing component contains motor preparatory information, we would further expect it to contain information regarding the parameters of the upcoming movement [Shenoy et al., 2013]. Indeed, the neural encoding weights in PMd (but not in M1; Supplementary Figure 6) were correlated across similar conditions and encoded both reach direction and curvature (Figure 2f-h). Therefore, while the trial-slicing components directly encoded motor sequences governing hand kinematics, the time-slicing component contained primarily preparatory information about movement parameters. Together, these results show that behaviorally relevant information in neural data can be spread across different slice types, motivating the need to demix variability classes with sliceTCA.

### 2.4 Pipeline for sliceTCA model selection and optimization

Dimensionality reduction methods, while powerful, can prove challenging in practice. First, robustly identifying the optimal number of components is a crucial yet challenging step in interpreting the dimensionality of neural representations [Stringer et al., 2019, Lanore et al., 2021]. In many tensor and matrix decomposition methods, such as NMF, different choices of the rank of the approximation may even lead to different results. Moreover, even after the rank is fixed, invariances in the decomposition may lead to multiple possible solutions. For example, matrix factorizations are known to be invariant to invertible linear transformations such as rotations. Similarly, sliceTCA is invariant to such transformations within each slice type (Supplementary Figure 7a, Methods). We have further identified a second class of invariant transformations that is specific to sliceTCA (Supplementary Figure 7b, 8, Methods). This invariance class, when unaccounted for, prevents an unambiguous attribution of covariance patterns to one of the three component types. Such invariances can lead to difficulties in comparing latent representations across multiple datasets [Williams et al., 2021] and are therefore crucial to address for any new method.

To address these concerns and to provide a user-friendly guideline for sliceTCA, we developed a full data analysis pipeline for sliceTCA including data preprocessing, model selection, model optimization, and visualization (Figure 3a). First, trials must be time-warped, trimmed, or masked, in order for the data to be shaped into a tensor. Second, for choosing the optimal rank, we developed a rigorous cross-validation procedure to identify the number of components of each slice type (Figure 3b), which we validated on ground-truth data (Supplementary Figure 9). Third, to address the invariances of the decomposition, we developed a hierarchical model optimization that adds additional constraints in the form of “sub-losses” that must be minimized at three stages of optimization (Figure 3c; Supplementary Figure 10). One of the stages of this procedure is a regularization of the reconstructed tensors of each slice type. Moreover, for non-positivity-constrained sliceTCA, the same criteria used for matrix factorization methods (e.g., maximization of variance and orthogonality as in PCA) can be applied to find unique solutions with respect to invertible linear transformations. We further prove mathematically that a unique solution is guaranteed if each of the sub-losses is unique (Supplementary Mathematical Notes). Together, employing a rigorous and standardized pipeline for model selection, fitting, and optimization allows the user to make a robust, principled choice of sliceTCA decomposition for further analyses and interpretation.

**Figure 3:**
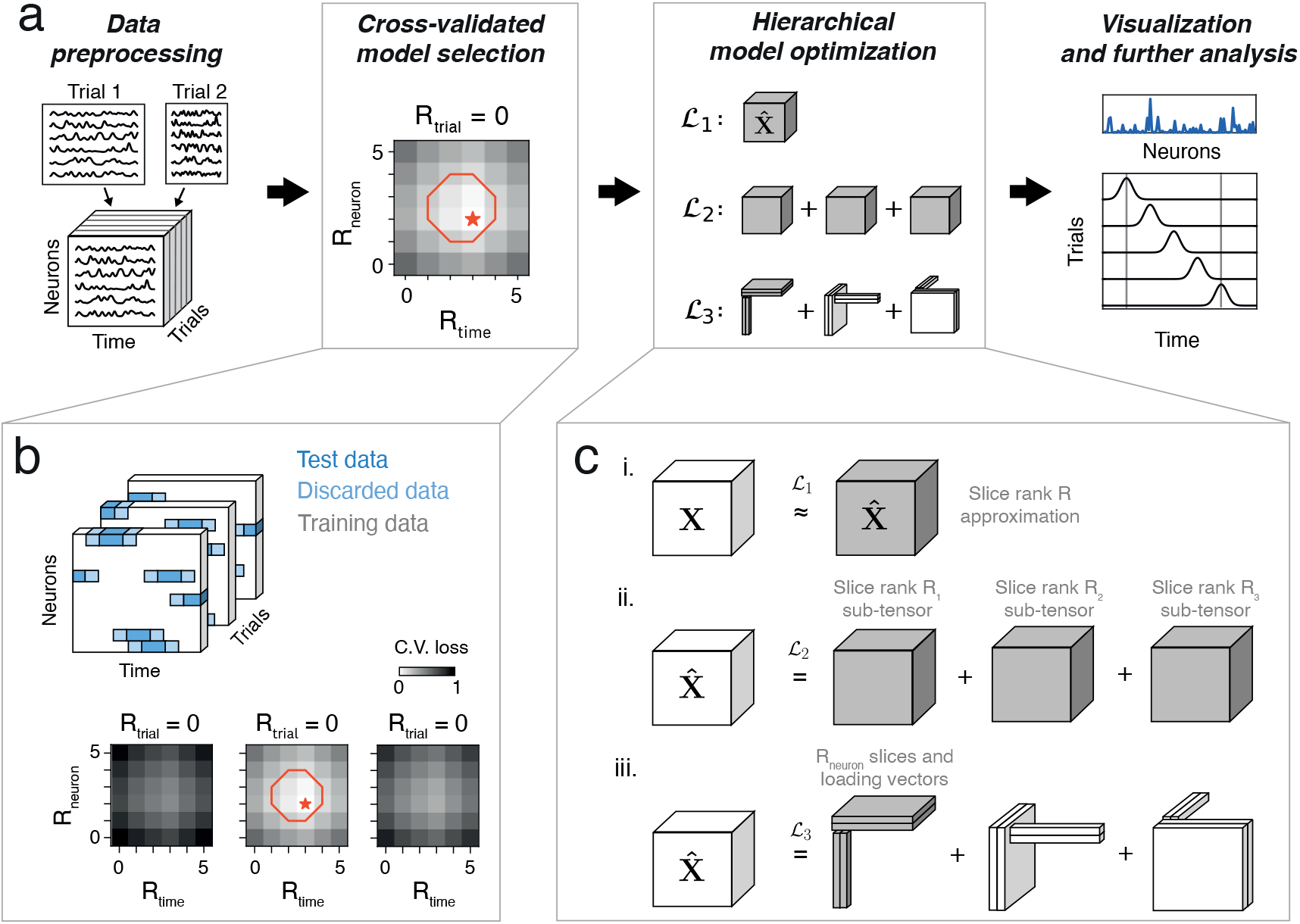
SliceTCA model selection, optimization, and analysis pipeline. **a.** SliceTCA data processing pipeline. First, neural data is preprocessed to form a data tensor. In experiments with variable trial length this could include temporal warping, exclusion of outlier trials, and/or trimming to the time period of interest. Second, model selection is performed to choose the number of components of each slice type (*R_neuron_, R_trial_, R_time_*) based on the cross-validated mean square error (MSE) loss. Next, the hierarchical model optimization procedure is performed to identify a unique decomposition for the model. **b.** The cross-validation procedure for neural data tensors that we propose. We randomly assign blocks of consecutive time points (blue) within the data tensor as held out data. The remaining entries of the tensor are used as training data (white). To reduce correlations between the training and testing data, we discard a brief period from the ends of the held-out blocks (light blue) from both training and testing. We use only the interiors of these blocks as test data (dark blue). We run a 3D grid search on the cross-validated loss (bottom). We either choose the optimal model (red star) or a model at the “elbow” of the loss function (red circle). **c.** Hierarchical optimization over the sliceTCA invariance classes. We first fit the model on all data (optimizing the MSE or 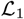 loss). Then, we consecutively optimize the secondary loss functions 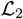 and 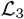 as described in (a). After this procedure, the resulting loading vectors and slices can be analyzed.

### 2.5 Denoising task-relevant manifolds during a motor task

With a standardized data analysis pipeline established, we next asked how behaviorally relevant latent structure sliceTCA could uncover in a novel dataset, without any prior expectation on the component types. We applied sliceTCA to a dataset consisting of simultaneously imaged cerebellar granule cells and pyramidal neurons in the premotor cortex of mice performing a motor task (Figure 4a) [Wagner et al., 2019]. Using the sliceTCA analysis pipeline, we selected a model with three trial-slicing components and three neuron-slicing components at the elbow of the cross-validated loss function (Figure 4b,c, Supplementary Figure 11; similar components observed in the optimal model, Supplementary Figure 12). The first trial-slicing component captured temporally distributed cerebellar and cortical dynamics that were common to both left and right correct reaches, but distinct from error reaches (Figure 4b,d). In contrast, the second trial-slicing component accounted for the differential activation in left vs. right trials (Figure 4b,d). A third component decayed slowly over trials, possibly representing adaptation over the course of the session (Figure 4b).

**Figure 4:**
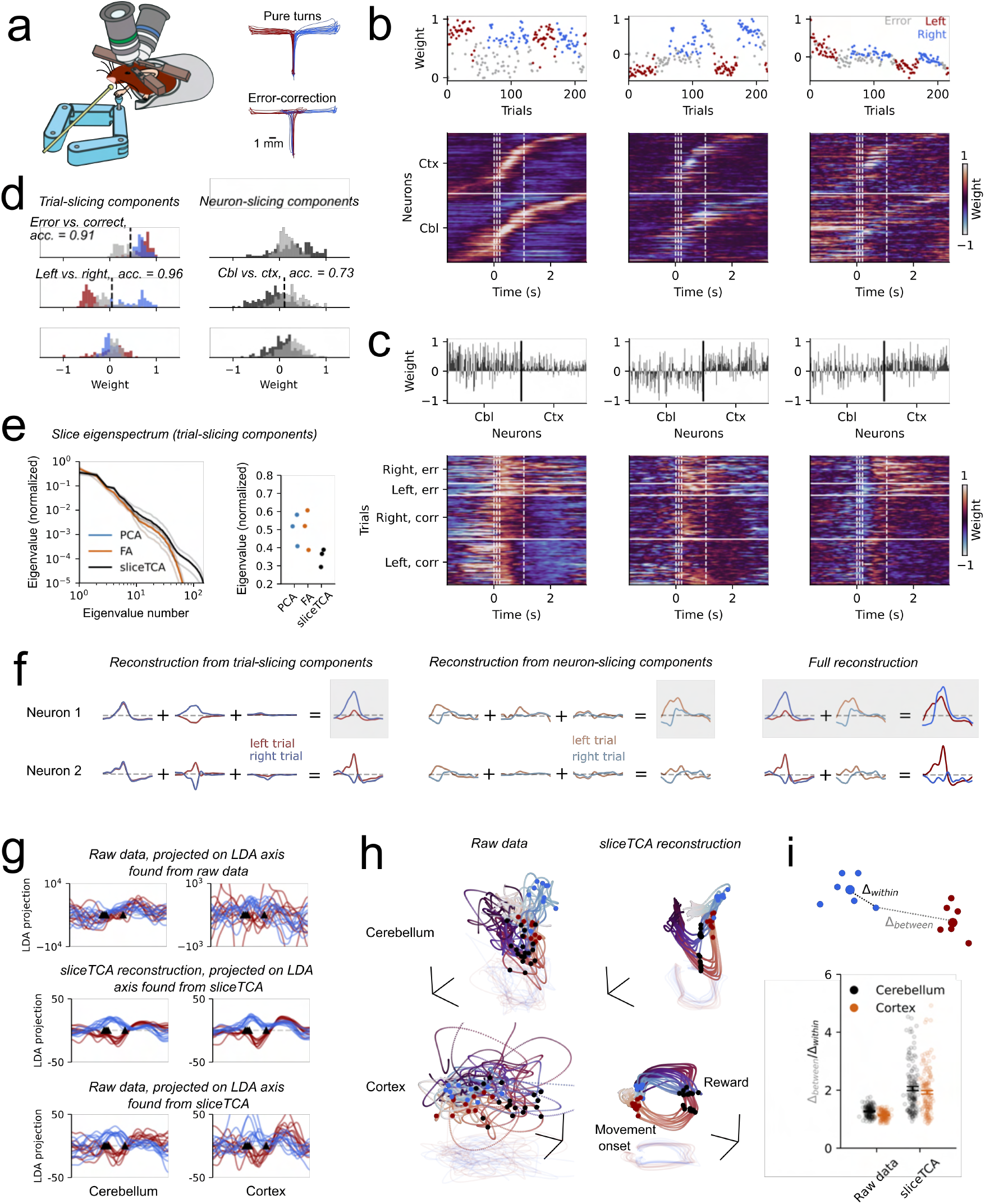
SliceTCA denoises task representations in simultaneously imaged cortical and cerebellar populations. **a.** Schematic of a mouse moving a manipulandum during simultaneous imaging of premotor cortex and cerebellum. Image modified from Wagner et al. [2019]. **b.** The three trial-slicing components identified by sliceTCA. Weights in the loading vectors are colored according to trial type: correct left (red), correct right (blue), and error (grey). In the slices, neurons are sorted within each region (cbl, cerebellum and ctx, motor cortex) by the latency of maximum activation in the first component. White dashed lines indicate movement onset, mid-turn, movement end, and reward. **c.** Three neuron-slicing components. Loading vectors are separated into cerebellar and cortical populations. In the corresponding slices, trials are separated into left or right cued trials and into correct or error turns (corr/err). Within each block, trials are plotted in increasing order (ascending). **d.** Histograms of loading weights for the three trial- (left) and neuron-slicing (right) components, colored by trial type and region. We classified weight vectors (correct vs. incorrect and left vs. right correct trials; cerebellum vs. cortex). **e.** To show that sliceTCA results in more demixed representations with higher-rank slices than concatenated matrix factorization methods, we calculated the eigenvalues of the slices of the three trial-slicing components identified by PCA, factor analysis (FA), or sliceTCA. Left: Slice eigenspectrum, averaged over the three trial-slicing components (black; spectra for individual components in grey). Right, leading eigenvalue for each component. **f.** Example reconstructions of low slice rank approximations of individual neurons. Left: Reconstruction from the trial-slicing components. The latent dynamics for each component are neuron-specific, but shared across trials up to a scaling factor. Middle: Reconstruction from the neuron-slicing components. The latent dynamics are trial-specific but shared across neurons except for a scaling factor. Right: The full sliceTCA reconstruction is obtained by summing the contributions of all components from both slice types. Red/blue indicate dynamics on an example left/right trial. **g.** Data from ten example trials per condition, projected onto an axis that maximally separates left and right correct trials between movement onset and reward. Upper, raw data projected onto an LDA dimension found from the raw data; middle, sliceTCA reconstruction projected onto a dimension found in the sliceTCA reconstruction; lower, raw data projected onto the LDA dimension found in the sliceTCA reconstruction. **h.** Neural manifolds comprising example trajectories per trial type in an orthonormalized neural subspace found with LDA (axis 1, same as g; axis 2 that separates activity at the time of movement onset vs. reward; axis 3 that separates pre-movement vs. mid-movement) from raw data and sliceTCA reconstruction. **i.** Separation of the left vs. right trajectories from full data and data denoised with a mixed-component sliceTCA model. Δ*_within_* (and Δ_*between*_) indicates the distance of the population vector in each trial around the time of movement onset to the center of the cluster of data points in its same (or, respectively, the opposite) trial class. Left and right trajectories are more separable after sliceTCA denoising (Wilcoxon signed-rank test, *p* < .001 both for cerebellum and motor cortex).

In addition, the three neuron-slicing components captured trial-specific population dynamics mostly localized around the time of movement or reward (dashed lines, Figure 4c), with prolonged activity in error trials, compared to correct trials, in the first and third component (Mann Whitney U-test, *p* < 0.001 for both components). Interestingly, the second neuron-slicing component captured differences between cerebellar and cortical activity (Figure 4c,d). The effect of simultaneously fitting two different covariability classes can be observed by comparing sliceTCA to matrix factorization methods that do not demix neural- and trial-covariability (Figure 4e, Supplementary Figure 13). While several loading vectors and slice weights of the components found by PCA and FA appear similar to their corresponding sliceTCA components, sliceTCA revealed more detailed structure for other components. But by contrast, without disentangling different covariability classes, the slices identified by PCA and FA were of lower rank than the sliceTCA slices (Figure 4e, Supplementary Figure 13), and thus capture less trial- or neuron-specific dynamics and less structure in the data. Together, these results show that sliceTCA identifies both task-specific (left, right, error trials) and region-specific (cerebellum vs. cortex) variables, by capturing the structure of neural data across multiple covariability classes.

Classic neural dimensionality reduction methods capture structure that is shared across neurons while removing variability that is specific to individual neurons. We next illustrate how additionally modeling structure that is neuron-specific but shared across trials affects the reconstruction of the data tensor (Figure 4f). Towards this end, we compared the neural representations of the raw data in neural space to the reconstructed data from the sliceTCA model. The sliceTCA reconstruction captured the same top principal components as the raw data, confirming that it was faithfully capturing the overall structure of the neural representation (Supplementary Figure 14). The advantage of including both neural and trial-covariability was reflected in increased behavioral interpretability of the neural representations. To show this, we projected the data onto the dimension that best separated left vs. right correct trials during the period between movement and reward. The axis found from the sliceTCA reconstruction revealed a more interpretable, denoised representation as compared to the dimension found from raw data (Figure 4g). Similarly, the task-relevant neural manifolds, found by projecting neural trajectories onto a subspace that separates activity along three task-relevant dimensions (see Methods), appear significantly denoised when sliceTCA was applied, compared to a direct projection of the raw data (Figure 4h; Supplementary Figure 14). We quantified this denoising effect by measuring the distance between left and right trials around the time of movement onset in sliceTCA reconstructions as compared to distances in raw data (Figure 4i). Our results show that sliceTCA, by grouping behaviorally similar trajectories in an unsupervised manner, increases the distance between trajectories of behaviorally distinct trials. Together, these results show that by demixing different classes of covariability, sliceTCA is able to denoise task-relevant representations in neural data in an unsupervised fashion.

### 2.6 Identifying components with region-specific covariability patterns in multiregion recordings

So far we have shown that mixed variability co-occurs within the same neural population. However, the need to consider multiple covariability classes becomes even more crucial in simultaneous recordings from many regions, as previous work has shown that different brain regions may be better described by different unfoldings of the data tensor [Seely et al., 2016]. Yet, relying on different tensor unfoldings for the analysis of distinct regions would require that these regions be analyzed separately, without leveraging the simultaneous nature of such data. We therefore asked whether sliceTCA could demix area-specific representations in distinct slice types.

To test this idea, we took advantage of a recently published dataset consisting of Neuropixel recordings across six brain regions during a perceptual decision-making task (Figure 5a) [IBL et al., 2022]. We selected a model with eight components: two trial-slicing components, three neuron-slicing components, and three time-slicing components (Supplementary Figure 15, 16). The two trial-slicing components identified variables related to behavioral performance (Figure 5b). The first trial-slicing component separated correct from incorrect trials (Mann Whitney-U test, *p* < .001), and the corresponding slice was characterized by reward-locked temporal response profiles in midbrain nuclei (APN and MRN), which we validated in single neuron PSTHs (Figure 5c). The second trial-slicing component instead featured more temporally heterogeneous responses in all regions and correlated inversely with log reaction times (Pearson’s *r* = −0.35, *p* < 0.001, *N* = 831 trials; Figure 5b). We next asked how these components contributed to the activity of different regions. The full sliceTCA reconstruction explained 33% −49% of neural activity, depending on the region (Figure 5d). Of this reconstructed activity, the two trial-slicing components contributed considerably to neurons in APN, MRN, and thalamus (TH) (19 ± 10%, mean ± s.d., *N* = 75 neurons; Figure 5e). Thus, the trial-slicing components identified stereotyped dynamics in subcortical regions TH, APN, and MRN that were linked to behavioral performance across trials.

**Figure 5:**
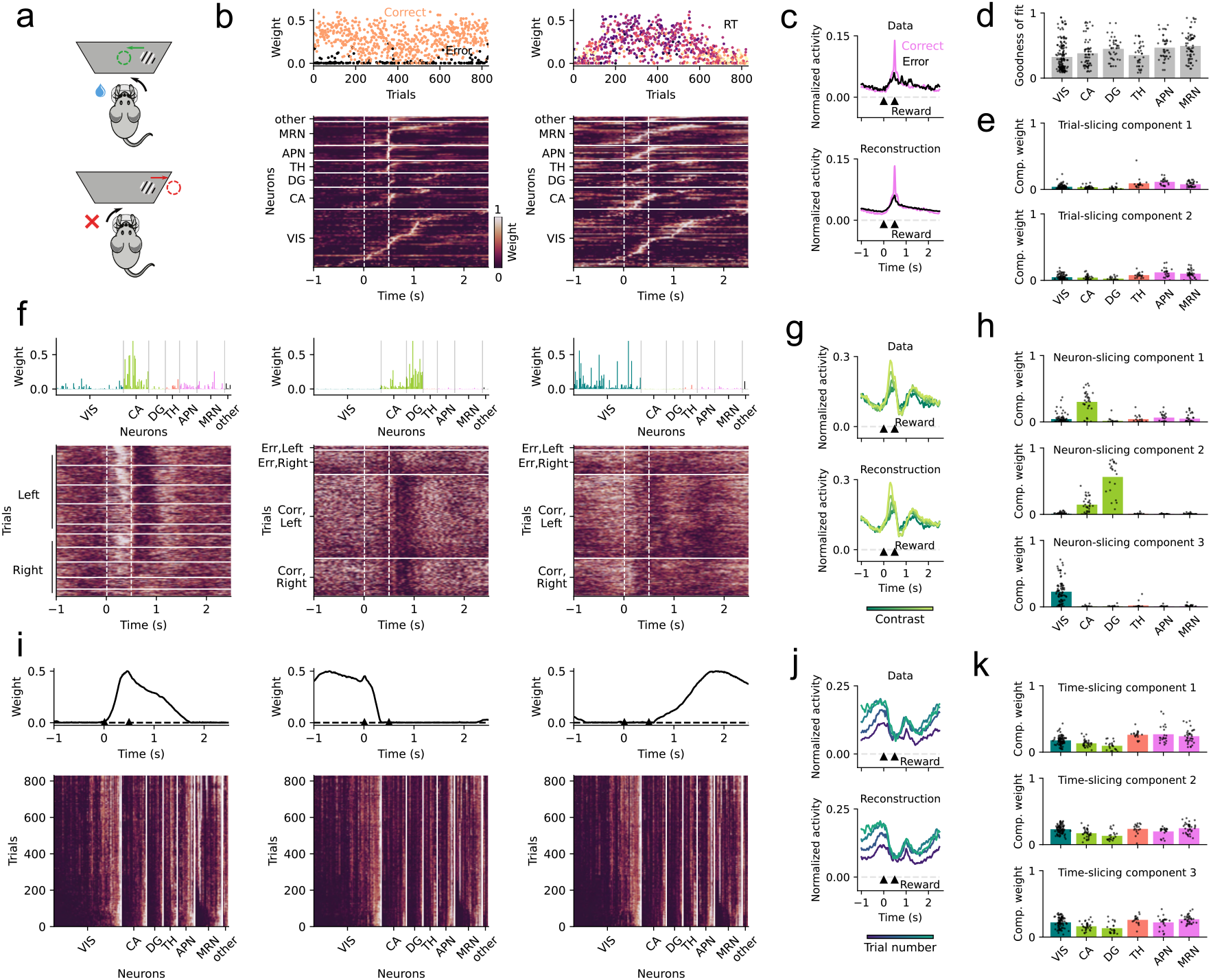
SliceTCA identifies region-specific sensory and behavioral variables in multi-region recordings. Schematic of perceptual decision making task from the International Brain Laboratory (IBL). Figure modified from IBL et al. [2021]. **b.** Two trial-slicing components: The loading vector of component 1 shows a separation between correct (orange) and error (black) trials. In component 2, the color scale in loading vector indicates log reaction time for each trial. In the corresponding slices: visual cortex (VIS), hippocampus (CA), dentate gyrus (DG), thalamus (TH), anterior pretectal nucleus (APN), and midbrain reticular nucleus (MRN). White dashed lines indicate stimulus onset and reward or timeout onset, respectively. Slice weights are normalized to [0,1] for each neuron separately and sorted by the latency of peak activation within each region (separately for each component). **c.** Top: PSTH of an example APN neuron with dominant reward-locked dynamics for correct (pink) and error trials (black). Bottom: PSTH built from the full sliceTCA reconstruction. Arrows indicate stimulus onset and reward. **d.** Reconstruction performance (Methods) of the full sliceTCA model, separated by region. Black dots indicate individual neurons. **e.** Contribution of each trial-slicing component to the overall reconstruction. **f.** Three neuron-slicing components: In each slice, trials are grouped into blocks separately for different components. In component 1 with dominant contribution of CA1, trials are grouped by contrast separately for left/right trials (within left/right, contrast increases from bottom to top). In components 2 (DG-related) and 3 (VIS-related), trials are grouped into blocks by left/right and correct/error. For all slices, within each block, trials are sorted in increasing order (ascending). Each slice is normalized to [0,1]. **g.** Top: PSTH of an example CA neuron for low to high contrasts (dark to light green). Bottom: PSTH built from the full sliceTCA reconstruction. **h.** Contribution of each neuron-slicing component to the overall reconstruction. **i.** Three time-slicing components: In the slices, neurons are sorted within each region according to increasing activation in early trials after normalizing weights for each neuron to [0,1] (same sorting across components). **j.** Top: PSTH of an example VIS neuron for early to late trials (blue to teal). Bottom: PSTH built from the full sliceTCA reconstruction. **k.** Contribution of each time-slicing component to the overall reconstruction.

In contrast, the three neuron-slicing components identified three distinct clusters of neurons corresponding to cortical regions: the hippocampus (CA), dentate gyrus (DG), and visual cortex (VIS) (Figure 5f). These components therefore represented population-wide covariability patterns that were specific to each of these regions. The slice of the CA-preferring component was characterized by a contrast-dependent activation between the sensory cue and reward (correlation of stimulus-evoked responses with contrast, Pearson’s *r* = 0.40, *p* < 0.001; Figure 5f,g), a feature which was less prominent in the DG and not observed in VIS-preferring components (*r* = 0.11, *p* = 0.002 for DG, *R* = −0.05, *p* = 0.14 for VIS). In the DG-preferring component, we observed post-reward suppression on correct (rewarded) trials which was significantly shorter on error trials (Mann Whitney U-test, *p* < 0.001; Figure 5f). The final VIS-preferring component revealed pre-stimulus activation that increased in strength over trials (Pearson’s *r* = 0.55, *p* < 0.001, Figure 5f), possibly indicating the emergence of a predictive signal of cue onset over the course of the experiment. Each component contributed to a large fraction of the sliceTCA reconstruction in its respective region (37 ± 21%, *N* = 138 neurons; Figure 5h). Therefore, the three neuron-slicing components represented different task-relevant features that were separately encoded in CA, DG, and VIS population responses.

Finally, the remaining time-slicing components partitioned the task duration into three distinct periods: early (pre-stimulus and stimulus onset), late (post-reward), and reward period (Figure 5i). The corresponding slices revealed smooth variations of the strength of each of these components in single neurons over the course of the experiment. Given the strong similarity of the three slices, we asked whether the components could sum to a flat trial-varying baseline for each neuron. However, when we examined single neurons we instead saw examples of a broad range of modulation patterns, with slowly varying activity that changed heterogenously over trials for the three task periods (Figure 5j as an example VIS neuron with increasing activity during pre-stimulus and post-reward periods, but not during the reward period). We tested this hypothesis using a linear model to compare the rate of change of the trial weights for each neuron across components (Methods). A substantial proportion of neurons across all regions showed significantly different rates of change across components (ANOVA, *p* < 0.05 with Bonferroni correction, *N* = 221 neurons; Supplementary Figure 17). Moreover, these three components contributed significantly to the sliceTCA reconstruction across all recorded regions (62±18%, *N* = 213 neurons; Figure 5h). Together, these results show that by accounting for different classes of covariability, sliceTCA is able to demix multi-region recording data into brain-wide representations of task period, and behaviorally-relevant stereotyped dynamics, and population-wide patterns of covariability encoded by individual regions.

### 2.7 Geometric interpretation of sliceTCA components

Recently, dimensionality reduction has been used in systems neuroscience to interpret neural population activity as trajectories embedded in a low-dimensional latent subspace within the full neural activity space. In sliceTCA, the neuron-slicing components can be interpreted in the same way due to their similarity to matrix factorization on trial-concatenated data. However, the time and trial slicing components have different interpretations as their natural bases lie within spaces in which each axis represents a different timepoint or a different trial. How then can then we grasp the time- and trial-slicing components’ contributions to latent representations in neural activity space?

We can answer this question by considering the hypothetical contribution from each slice type separately. First, note that while the neuron-slicing components are constrained to an *R*_neuron_-dimensional subspace, their dynamics within that subspace are unconstrained over trials (Figure 6, neuron-slicing component). On the other hand, the dynamics of the *R*_time_ time-slicing components are constrained to a common temporal dynamic, but the neural weight vectors can instead vary from trial to trial. Geometrically, this means that the reconstruction from these components lies within an *R*_time_-dimensional subspace that can now vary on each trial, but that the temporal dynamics within each trial-specific subspace is constrained to be the same (Figure 6, time-slicing component). Finally, the *R*_trial_ trial-slicing components’ neural weights change at every timepoint, while trial weights are fixed. This corresponds to latent dynamics that are no longer embedded in a low-dimensional subspace, but that are built instead from stereotyped dynamical trajectories (Figure 6, trial-slicing component; but see Supplementary Figure 18). In this way, the three covariability classes that we have described can also be seen as three classes of latent dynamics in neural activity space. Together, all three classes contribute to the dynamics of the full reconstruction, which may appear more complex than any one component type (Figure 6a, reconstruction).

**Figure 6:**
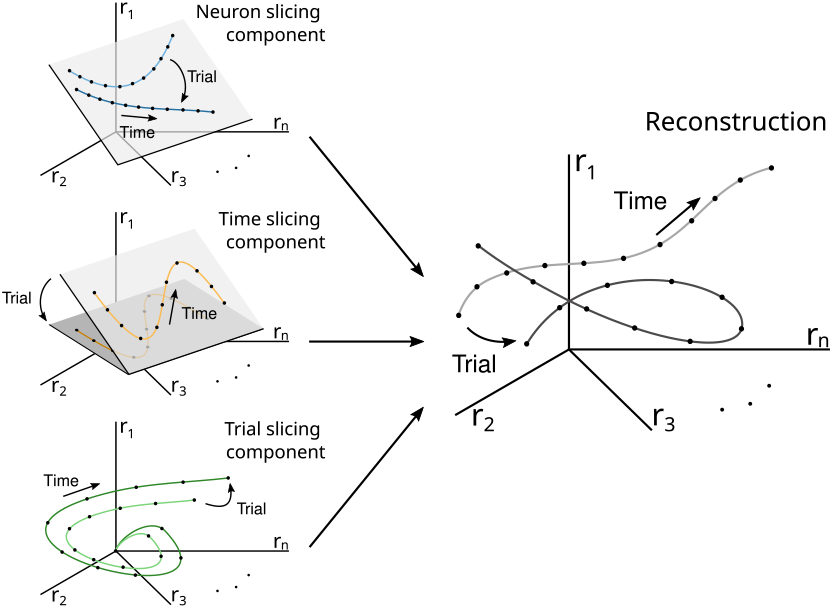
Different slice types capture latent variables with distinct geometric properties. *Neuron-slicing component.* Example of two neuron-slicing components visualized in neural activity space. The latent trajectories are embedded in an two-dimensional subspace, but their dynamics within that subspace are unconstrained. *Time-slicing component.* Example of two time-slicing components. These are similarly embedded within an twodimensional subspace, but that subspace varies over trials. The latent variables are further constrained to follow the same dynamics within each latent subspace. *Trial-slicing component.* The trial-slicing components are not constrained to any latent subspace, as the neural encodings may change at every timepoint. These components describe potentially high-dimensional dynamics that are stereotyped across trials. Note that here only 1 component is shown for clarity. *Reconstruction.* After summing these components, the full latent trajectories are not necessarily limited by any of the geometric constraints that characterize individual slice types.

This geometric view illustrates that by fitting different classes of covariability, sliceTCA is able to capture latent dynamics that are no longer confined to a linear subspace, despite still being a multilinear method. In contrast, traditional matrix factorization methods which capture only a single covariability class are restricted to one of the three geometric classes of latent dynamics in neural space shown in Figure 6, while TCA constrains its components to obey the geometrical constraints of all three classes simultaneously (Supplementary Figure 19). In sum, sliceTCA is able to capture a broader range of covariability structure in neural data, and a broader range of latent representations in neural space, than related methods, all while remaining easily interpretable.

## 3 Discussion

Neural population dynamics are frequently interpreted as low-dimensional latent variables encoded by fixed subgroups of neurons, which represent shared variability across neurons. Here, we have advocated for an expansion of this view of structure in neural data which takes into account three distinct classes of shared variability: across neurons, time, and trials. Towards this end, we introduced sliceTCA, a new tensor decomposition method that is able to demix latent variables that belong to any of these covariability classes. Through several example datasets, we demonstrated that sliceTCA can capture more task-relevant covariability in neural data in fewer components, enabling the description and interpretation of complex latent structure embedded in large-scale neural recordings. Finally, we illustrated how sliceTCA expands the classic view of neural population dynamics towards latent variables that are not constrained to low-dimensional dynamics in a fixed, linear manifold.

Our framework of multiple covariability classes addresses key limitations of the classic view on latent dynamics, which is unable to identify several types of structure commonly found in neural data. In particular, this view fails to capture neural sequences, as previously pointed out in the literature [Mackevicius et al., 2019, Seely et al., 2016]. Indeed, task-relevant neural sequences are a widespread phenomenon observed across brain regions during navigation, timing, value-based decision making, and motor production [Harvey et al., 2012, Parker et al., 2022, Zhou et al., 2020]. Here we have emphasized the ability of the trial covariability class to capture neural sequences that have shared structure across trials, e.g. choice-specific sequences. However, we note that this class can capture more complex forms of neuron-specific temporal patterning within a trial [Feng et al., 2015, Lakshmanan et al., 2015, Koay et al., 2022]. On the other hand, population modes characterized by trial-to-trial differences in timing are captured in the neural covariability class. Such variations in timing may be critical for interpretation, for example the shift of the reward prediction error during temporal difference learning [Amo et al., 2022, Schultz, 1998]. Lastly, the time covariability class may be well-suited for describing forms of learning or representational drift in which the latent space over which neural data evolves over trials [Hennig et al., 2021, Rule et al., 2019]. Importantly, it has been argued that different brain regions are better described by neural or by trial covariability [Seely et al., 2016]. Our results support this hypothesis, and further show that these different classes can be demixed by sliceTCA. Therefore, demixing covariability classes may be a crucial step when considering large-scale multi-region recordings [IBL et al., 2022, Wagner et al., 2019, Ebrahimi et al., 2022, Ahrens et al., 2012].

A longstanding challenge in systems neuroscience is the difficulty of mapping neural variability to changes in behavior [Renart and Machens, 2014]. This can be accomplished using supervised dimensionality reduction methods that use information regarding behavior or task outcomes to identify latent variables [Kobak et al., 2016, Sani et al., 2021a, Balzani et al., 2022]. Despite it being an unsupervised method, we found that SliceTCA was able to disentangle behavioral and task information in each of the datasets presented. We claim that this is due to two reasons: first, demixing different sources of covariability effectively “denoises” components that represent task variables that would have otherwise been occluded by additional sources of variability. Second, the trial-slicing components explicitly identify dynamics that are shared across trials, which tend to be defined by task variables or behavioral outcomes. Indeed, in each of the three datasets, we found that trial-slicing and time-slicing components correlated with behavioral variables. Moreover, in our feedforward model, we suggest how sliceTCA could offer a window into the computational roles of variables modeled by different slice types. We argue that the classical view on neural latents, which assumes that a key part of behaviorally relevant neural variability is correlated across neurons, is overly reductionist and may miss many types of neural dynamics underlying behavior.

Beyond tensor and matrix based methods, more sophisticated forms of nonlinear dimensionality reduction can be used to identify latent variables embedded within a curved manifold [Balasubramanian and Schwartz, 2002, Belkin and Niyogi, 2003, McInnes et al., 2018]. Within neuroscience, several methods have been proposed specifically for neural data, including methods based on neural networks [Pandarinath et al., 2018, Schimel et al., 2022, Sani et al., 2021b] or manifold reconstruction based on topological features [Chaudhuri et al., 2019, Rybakken et al., 2019]. While these methods are crucial for identifying nonlinearly embedded latent variables, a key advantage of matrix and tensor decompositions is the simplicity of the models. Indeed, the analytical tractability of the sliceTCA decomposition enabled us to characterize its invariance classes and to propose a method to identify a unique solution. Identifying invariances is crucial for reproducibility and interpretation, as non-unique solutions may prohibit clear comparison across datasets [Dyer et al., 2017, Gallego et al., 2020]. This issue is ever more important with the recent increase in popularity of comparisons of neural data to task-trained neural network models, whose representations are known to be sensitive to model specifications such as architecture and inputs [Lindsay et al., 2022, Williams et al., 2021]. Going forward, matrix and tensor decompositions could prove useful for comparing latent representations by virtue of their interpretability and tractability.

SliceTCA falls into a larger class of tensor decomposition methods including TCA [Williams et al., 2018, Harshman et al., 1970], which captures variability lying at the intersection of the covariabilty classes, and the Tucker decomposition, which allows factors to interact via a core tensor [Onken et al., 2016]. Yet while tensor decompositions can be viewed as generalizations of matrix factorizations, they do not always have the same properties. For example, tensor-based methods are known to be generally more computationally expensive [Kolda and Bader, 2009, Bläser et al., 2019]. Still, tensor decompositions are key methods in neuroscience as they allow the discovery of components that can be mapped across trials or conditions. Here we have focused on the classic third-order tensors (neurons × time × trials) that are frequently used in neuroscience. Current experimental techniques are rapidly enabling the acquisition of data tensors of even higher order, by adding legs that correspond to days or conditions. Future extensions of tensor methods that allow individuals to be incorporated as an additional leg could help to identify the neural basis of variability across subjects [Kuchibhotla et al., 2019, Smith et al., 2022]. Going forward, our framework of mixed classes of covariability can help to advance our understanding of behaviorally relevant latent structure in high-dimensional neural data recorded during increasingly complex tasks, across brain regions and across individual subjects.

## 4 Methods

### 4.1 Definition of sliceTCA model

#### 4.1.1 Matrix rank and matrix factorization

Consider a data matrix consisting of *N* neurons recorded over *T* samples (timepoints): **X** ∈ ℝ^*N×T*^. Matrix factorization methods find a low-rank approximation 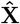 following Eq. 1, in which each component is a rank-1 matrix: **X**^(*r*)^ = **u**^(*r*)^ ⊗ **v**^(*r*)^, where **u**^(*r*)^ ∈ ℝ^*N*^ and **v**^(*r*)^ ∈ ℝ^*T*^ are vectors representing the neural and temporal coefficients, which are chosen to minimize a loss function. In other words, the activity of neuron *n* at time *t* is given by:

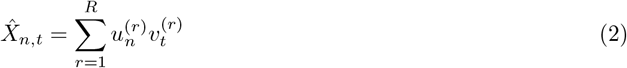

A common choice of loss function is the mean squared error:

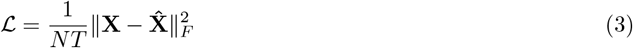

Constraints may be added to the minimization of the loss, such as non-negativity of the coefficients in NMF.

#### 4.1.2 Slice rank and sliceTCA

A d-tensor is a generalization of data matrices to *d* legs (i.e, a data matrix is a 2-tensor). Here we are specifically concerned with 3-tensors typically used in neuroscience, in which the three legs represent neurons, time, and trial/condition: **X** ∈ ℝ^*N*×*T*×*K*^. SliceTCA extends the matrix factorization in Eq. (1) by fitting **X** with a low *slice rank* approximation [Tao and Sawin, 2016]. A slice-rank-1 d-tensor is an outer product of a vector and a (*d* – 1)-tensor. For the 3-tensors that we have been considering, this corresponds to the outer product of a ‘loading’ vector and a 2-tensor, thus making this 2-tensor a *slice* of this slice-rank-1 tensor up to a scalar multiple determined by the loading vector.

Each sliceTCA component can be one of three different slice types. For example, a neuron-slicing component can be written as **X**^(*r*)^ = **u**^(*r*)^ ⊗ **A**^(*r*)^ where **A**^(*r*)^ ∈ ℝ^*T*×*K*^ is the time-by-trial slice representing the dynamics of the component across both time and trials and the vector **u**^(*r*)^ represents the neural loading vector. Components of other slice types can be constructed similarly with their respective loading vectors and slices: **v**^(*r*)^ ∈ ℝ^*T*^, **B**^(*r*)^ ∈ ℝ^*N*×*K*^ for the time-slicing components, and **v**^(*r*)^ ∈ ℝ^*K*^, **C**^(*r*)^ ∈ ℝ^*N×T*^ for the trial-slicing components. Put together, this results in a decomposition of the following form:

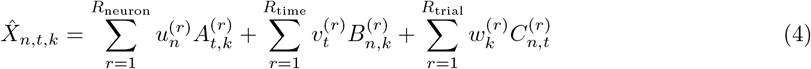

Because of the different slice types, each sliceTCA model can be described by the hyperparameter 3-tuple **R** = (*R*_neuron_,*R*_trial_,*R*_time_), defining the number of neuron-, trial-, and time-slicing components, for a total of *R*_neuron_ + *R*_trial_ + *R*_time_ components.

#### 4.1.3 Relationship to TCA

The extension of matrix factorizations to TCA is based on a different definition of tensor rank, in which a rank-1 tensor is as an outer product of d vectors. Each component is defined by a set of vectors corresponding to trial coefficients **w**^(*r*)^ ∈ ℝ^*K*^ to each component: **X**^(*r*)^ = **u**^(*r*)^ ® **v**^(*r*)^ ® **w**^(*r*)^. Then each element of the approximated data tensor can be written as:

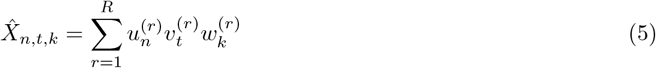

In other words, a TCA component is a special case of a sliceTCA component in which the slice is a rank-1 matrix. In this way, sliceTCA is more flexible than TCA as it has fewer constraints on the type of structure that is identified in the data. However, this increase in flexibility comes with a cost of an increase in the number of parameters, as sliceTCA fits all the entries of each slice. The flexibility of sliceTCA also leads to different invariance classes as discussed below. Finally, we note that the two methods can in principle be merged by incorporating TCA components into Eq. 4.

### 4.2 SliceTCA invariance classes

#### 4.2.1 Transformations within a slice type

Matrix factorization methods are known to be invariant to invertible linear transformations, including, but not limited to, rotations of the loading vectors. For example, suppose we decompose a matrix **Y** ∈ ℝ^*N×T*^ into a the product of a matrix of weights, **W** ∈ ℝ^*N×R*^ and a matrix of scores, **S** ∈ ℝ^*R×T*^. Consider any invertible linear transformation **F** ∈ ℝ^*R×R*^. Then **Y** can be re-written as:

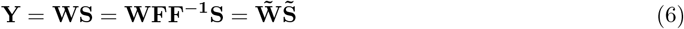

where 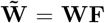 and 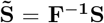. As a result, matrix decompositions like factor analysis (FA) lead to not one solution, but rather an invariance class of equivalent solutions. Note that PCA avoids this problem by aligning the first component to the direction of maximum projected variance, as long as the eigenvalues of the covariance matrix are distinct. However, other methods which do not have a ranking of components are not able to use the same alignment. SliceTCA inherits this same invariance class, since all the loading vectors within a given slice type can be transformed in the same way as Eq. (6) to yield the same partially reconstructed tensor for each slice type (Supplementary Figure 7a).

#### 4.2.2 Transformations between slice types

SliceTCA has an additional invariance class due to the fundamental properties of multilinear addition. For example, consider a slice-rank-2 tensor **Y** ∈ ℝ^*N×T×K*^ which is made of two components of different slice types, which we will assume without loss of generality to be neuron- and time-slicing components with corresponding slices **V** and **U**, such that:

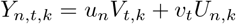

Then the following transformation can be performed for arbitrary vector **z** ∈ ℝ^*K*^,

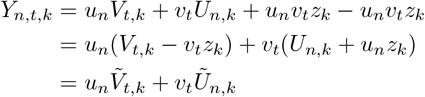

where 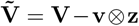 and 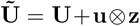 are transformations of the original slices. This invariance class therefore corresponds to passing a tensor-rank-1 tensor between two slices of differing slice types (Supplementary Figure 7b).

Note that two classes of transformations (within-slice-type and between-slice-type) commute (see proposition 1.2 of Mathematical Notes), and therefore one cannot get a new transformation by, for example, applying the first transformation, the second, and then the first again.

#### 4.2.3 Identification of unique sliceTCA decomposition

In order to find a uniquely defined solution we can take advantage of natural hierarchy between the two invariance classes. Specifically, let us first define the partial reconstruction 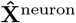 of the low-slice-rank approximation 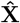 based on the neuron-slicing components, i.e.:

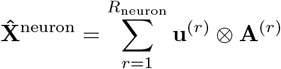

and let 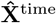 and 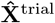 be similarly defined, so that 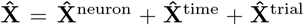. Now note that the within-slice-type transformations change the weights of the loading vectors and slices of all components of a given slice type, without changing the partial reconstructions for each slice type. For example, applying these transformations to the neuron-slicing components would change **u**^(*r*)^ and **A**^(*r*)^ but not 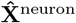. On the other hand, the between-slice-type transformations change the partial reconstructions 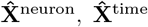 and 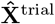, but not the full reconstruction 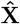.

We leveraged this hierarchy to develop a post-hoc model optimization into three steps, each with a distinct loss function. The first step identifies a model that minimizes a loss function 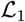 defined on the full reconstruction (Figure 3c i), fixing the tensor approximation 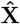. Next, we use stochastic gradient descent to identify the between-slice-type transformation that minimizes a new loss function 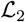, which fixes 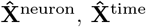 and 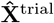 without affecting 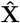 (Figure 3c ii). Finally, we identify the within-slice-type transformation that minimizes loss 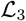 to arrive at the final components (loading vectors **u**^(*r*)^, **v**^(*r*)^, **w**^(*r*)^ and slices **A**^(*r*)^, **B**^(*r*)^, **C**^(*r*)^) without affecting 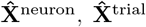, and 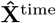 (Figure 3c iii). Each of the three loss functions can in principle be chosen according to the constraints or normative assumptions most relevant to the question at hand. Furthermore, we prove that if each of these objective functions leads to a unique solution, then the decomposition is unique under the condition that rank 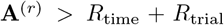 for all *r* = 1,..., *R*_neuron_ and similarly for the other two slice types (see Theorem 1.8, Supplementary Mathematical Notes).

### 4.3 Model selection, optimization and fitting

To fit sliceTCA for a given dataset arranged as a 3-tensor, we followed the data analysis pipeline described in the main text. Below, we provide details and hyperparameters for the steps involved in the pipeline.

#### 4.3.1 Fitting sliceTCA with stochastic gradient descent

For a fixed choice of **R**, model parameters (i.e., loading vectors and slices for each component) were fitted using the optimizer Adam [Kingma and Ba, 2014] in Pytorch [Paszke et al., 2019]. Initial parameters were randomly drawn from a uniform distribution over [-1, 1] or [0, 1], respectively, for unconstrained and non-negative sliceTCA. Throughout, we optimized the mean-squared error (MSE) loss in Eq. (3) with a learning rate of 0.02. To introduce stochasticity in the computation of the gradient, and thus avoid local minima, we masked a fraction of tensor entries so that they are not included in the calculation of the loss. This fraction starts at 80 % and decreases exponentially during training with a decay factor of 0.5 over three (Figure 2) or five blocks of iterations (Figures 4 and 5), respectively. Within each block, the mask indices are randomly reinitialized every 20 out of a total of 150 (Figure 2), 200 (Figure 4), or 100 iterations per block (Figure 5). To obtain an optimal model under a given **R**, we repeated the fitting procedure ten times with different random seeds and chose the model with the lowest loss.

#### 4.3.2 Cross-validated grid search

To choose the number of components in each slice type, we run a three-dimensional grid search to optimize the cross-validated loss. In addition to the decaying mask used during model fitting, we mask 20 % of the entries throughout the fitting procedure as held-out data. These masked entries were chosen in randomly selected 1 s (Figure 4) or 150 ms blocks (Figure 5) of consecutive timepoints in random neurons and trials. Blocked masking of held-out data (rather than salt-and-pepper masking) was necessary to avoid temporal correlations between the training and testing data due to the slow timescale of the Ca^2+^ indicator or due to smoothing effects in electrophysiological data. To further protect against spuriously high cross-validation performance due to temporal correlations, we trimmed the first and last 250 ms (Figure 4) or 40 ms (Figure 5) from each block; this data was discarded from the test set, and only the remaining interior of each block was used to calculate the cross-validated loss. We repeated the grid search ten times with different random seeds for train-test-split and parameter initialization, while keeping a constant seed for different **R**. Once the cross-validated grid search is complete, we selected **R**\* by identifying the model with minimum or alternatively, near-optimal average test loss across seeds. Admissible models are defined as achieving a minimum of 80 % of the optimal performance for non-constrained sliceTCA, and 95 % of the optimal model performance for non-negative sliceTCA, as compared to root mean squared entries of the raw data.

#### 4.3.3 Hierarchical model optimization

For the first step of the model optimization procedure, we chose the mean squared error loss for 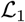:

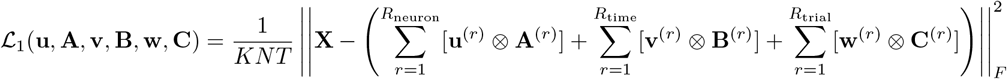

as in the model selection (essentially refitting the model with the specific ranks identified with the cross-validation procedure on the entire data). For 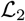 we use the sum of the squared entries of the three partial reconstructions from each slice type,

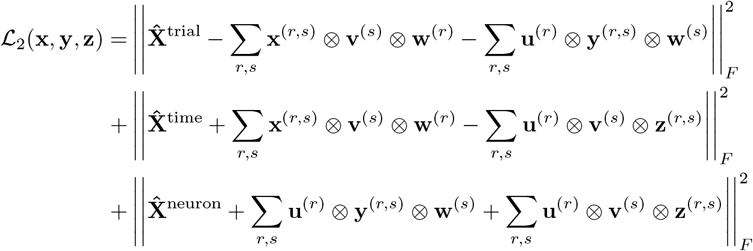

where 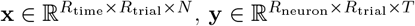, and 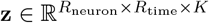. This can be thought as a form of L2 regularization. For 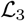 we chose orthogonalization and variance explained ordering through singular value decomposition.

### 4.4 Feedforward model of perceptual learning

We modeled a population of linear neurons receiving sensory input from upstream sources representing a Go and a No-go stimulus, as well as input representing top-down modulation which varied from trial to trial. On each trial *k*, either the Go or No-go stimulus was activated, with probability p = 0.5 of presenting the same stimulus as was presented in the previous trial. Go/No-go inputs *x^GO^,x^NO^* were assumed to follow the same bell-shaped activation function *s_t_* = *e*^-(*t*-4)^ on the trials during which their corresponding stimulus was presented, i.e., 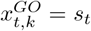 if *k* was a GO trial, 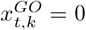 otherwise (and vice versa for No-go input).

The stochastic learning process of the Go and No-go weights 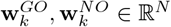 over trials was modeled as a Ornstein-Uhlenbeck process, which was initialized at 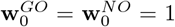 and evolved independently across neurons:

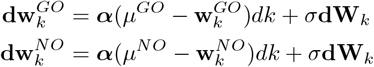

where 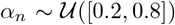 are the neuron-specific learning rates, *μ^GO^* = 2, *μ^NO^* = 0, *σ* = 1.3. Furthermore, to keep weights non-negative and simulate their saturation, they were clamped to [0, 2]. The process was evaluated using a stochastic differential equation solver and sampled at *K* evenly spaced points in [0,10] representing *K* trials.

Top-down modulation was modeled as a rectified Gaussian Process:

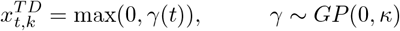

with temporal kernel:

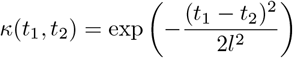

where 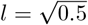. Top-down weights were non-plastic and distributed as 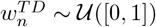. The activity of each neuron was thus given by:

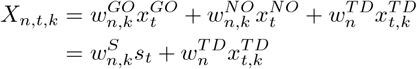

where the sensory input is combined into 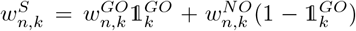 where 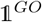 is an indicator function that is 1 when trial *k* is a Go trial and 0 if it is a No-go trial. By construction, the tensor **X** has slice rank of 2, as it can be written in the following form:

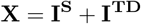

where 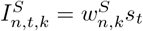 is a time-slicing component representing the weighted, trial-specific sensory input and 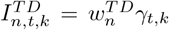 is a neuron-slicing component representing top-down modulatory factors that vary over trials. In our simulations, we used *K* = 100, *T* = 90, *N* = 80.

We fitted sliceTCA with non-negativity constraints to the synthetic dataset, using five blocks of 200 iterations each with a learning rate which decayed exponentially over blocks from 0.2 to 0.0125, and a mask that decayed exponentially over blocks from 0.8 to 0.05. Masked entries changed randomly every iteration. Initial parameters were drawn uniformly over [0, 1].

### 4.5 Dataset 1: Motor cortical recordings during a center-out and maze reaching task

#### 4.5.1 Description of the dataset

We analyzed a dataset of motor cortical (M1, *N* = 90) and premotor cortical electrophysiological recordings (PMd, *N* = 92) [Churchland et al., 2012], which is curated and publicly available as part of the ‘Neural Latents Benchmark’ project [Pei et al., 2021]. Briefly, monkeys were trained to perform a delayed center-out reach task to one of 27 locations in both maze conditions (in which barriers were placed on the screen, leading to curved optimal reach trajectories) and in no maze conditions with matched target locations (classic center-out task leading to straight optimal reach trajectories). The go signal for movement initiation appeared 0 — 1000ms after target onset and 1000 — 2600ms after the trial started with a fixation cue. We analyzed data from one animal (monkey J) in a single session and randomly subselected 12 target locations, resulting in *K* = 246 single-target trials in the maze reach conditions and *K* = 265 single-target trials in the 12 center-out reach conditions with matched target locations.

#### 4.5.2 Additional preprocessing

We calculated firing rates for bins of 10 ms which we then smoothed with a Gaussian filter with *σ* = 20 ms and rescaled to minimum and maximum values of 0 and 1 over the course of the experiment for each neuron separately. We selected a time period starting 1 s before movement onset (thus including a substantial part the motor preparation period) and ending 0.5 s after movement onset, when the monkey had successfully reached the target position in the majority of trials. We did not time-warp the data. The resulting data tensor had dimensions of *N* = 182, *T* = 150, and *K* = 511.

#### 4.5.3 Supervised mapping of neural population activity onto kinematic data

To identify the neural subspace from which 2D hand trajectories could be read out (Figure 2a), we used ordinary least squares (OLS). Specifically, we found weights that project the neuron-unfolded data from the full neural space onto a 2D subspace that best maps onto x/y hand velocity with a time delay of 100 ms to account for the lag between neural activity and movement. When testing the decoding analysis after dimensionality reduction we instead applied OLS to the reconstruction (or partial reconstruction, i.e., from only a single slice type) after reshaping it into a *N* × *KT* matrix. We also used OLS to project time-averaged pre-movement activity onto target locations (Figure 2g). For Figure 2h, we used LDA to identify the dimension that best separates pre-movement averaged activity in clockwise vs. counter-clockwise curved reaches in the maze condition. To plot activity in a 3D neural subspace that contained information about the upcoming movement, we then orthogonalized the two axes which map neural activity onto target locations to the axis that distinguishes clockwise and counter-clockwise movements.

For all decoding analyses, we calculated *R*^2^ values on left-out trials in a 5-fold cross-validation procedure performed on 100 permutations of the trials. Decoding was performed on data from period spanning 250 ms before to 450 ms after movement onset. For trial-resolved data (Figure 2a, raw data, neuron-slicing, TCA, trial-slicing.), we averaged trial-wise *R*^2^ values, and for pre-movement information on target positions, we calculated a single *R*^2^ value across trials for center-out and maze reaching conditions. For trial-averaged data (Figure 2a, trial-averaged), we performed 2-fold cross-validation by averaging hand and neural trajectories separately for each fold, and then calculating *R*^2^ values averaged over conditions and folds.

#### 4.5.4 Visualization of sliceTCA weights

The results of fitting non-negative sliceTCA are shown in Figure 2c,d and Supplementary Figure 3. Each component consists of a weight vector and a slice of corresponding weights on the other two variables. Along the trial dimension, we sorted trials by the angle of the target position and whether trials belonged to center-out or maze reaching conditions. Along the neuron dimension of trial-slicing components, neurons were sorted by the peak latency of neural activity in the first component. For the time-slicing component, neurons were sorted according to their mean activity in the first reaching condition.

#### 4.5.5 Correlation matrices

To assess the encoding similarity of movement preparation in the time-slicing component, we calculated the *K* × *K* correlation matrix of the neural encoding weights (i.e., the rows of the slice in Figure 2d) for different pairs of trials, separately for center-out and maze reach conditions, and for PMd (Figure 2f) and M1 (Supplementary Figure 6). We sorted the resulting correlation matrices by the angle of the target location (Figure 2f).

### 4.6 Dataset 2: Cortico-cerebellar calcium imaging during a motor task

#### 4.6.1 Description of the dataset

We analyzed recently published calcium imaging data consisting of simultaneously recorded cerebellar granule cells (*N* = 134) and premotor cortical L5 pyramidal cells (*N* = 152) from a head-fixed mouse performing a motor task in which a manipulandum had to be moved forward and left- or rightward for a reward [Wagner et al., 2019]. After a correct movement was completed, a water reward was delivered with a 1 s delay, followed by an additional 3.5 s inter-trial interval. Left vs. right rewarded turn directions were alternated without a cue after 40 successful trials. We analyzed data from one sessopm of a mouse in an advanced stage of learning, comprising a total of *K* = 218 trials. The data was sampled at a 30 Hz frame rate. Calcium traces were corrected for slow drifts, z-scored and low-pass filtered [Wagner et al., 2019].

#### 4.6.2 Additional preprocessing

Due to the freely timed movement period, we piecewise linearly warped data to the median interval lengths between movement onset, turn, and movement end, respectively. The remaining trial periods were left unwarped and cut to include data from 1.5 s before movement onset until 2.5 s after reward delivery, resulting in a preprocessed *N* × *T* × *K* data tensor with *N* = 286, *T* = 150, and *K* = 218.

#### 4.6.3 Visualization of sliceTCA weights

In Figure Figure 4b,c, we show the results of a fitted sliceTCA model. We further reordered trials in the trial-time slices according to trial type, and the neurons in neuron-time slices according to the peak activity in the first trial-loading component. This allows for a visual comparison of tiling structure across components. We used Mann Whitney U-tests on time-averaged activity between reward and end of trial in trial-time slices. We used LDA to determine the classification accuracy for neuron identity (cerebellum vs. cortex) based on the loading vector weights of the three neuron-slicing components found by sliceTCA. We similarly reported classification accuracy of trial identity (error vs. correct, left vs. right) based on the loading vector weights of the trial-slicing components.

#### 4.6.4 Matrix rank of slices

To determine whether sliceTCA finds components with higher matrix rank than methods that do not demix slice types (neuron-slicing PCA and factor analysis (FA) with neuron loadings, neuron-time-concatenated PCA and FA with trial loadings), we performed singular value decomposition (SVD) on the six slices of the sliceTCA model shown in Figure Figure 4b, as well as on the scores of either trial-slicing or neuron-slicing PCA and FA, after refolding the resulting scores into *N* × *T* or *K* × *T* matrices, respectively. We then compare these to the normalized eigenvalue spectra of the slices of the trial-slicing (Figure 4e) or neuron-slicing components (Supplementary Figure 13d). Factor analysis was performed using the Python package “sklearn” [Pedregosa et al., 2011], which uses an SVD-based solver. For comparability with PCA and sliceTCA solutions, no factor rotations were performed.

#### 4.6.5 Manifolds from sliceTCA reconstructions

To analyze the geometry of neural data, we reconstructed the low-slice-rank approximation of neural activity from the sliceTCA model separately for the cerebellum and for the premotor cortex. We then used LDA on both raw and reconstructed data to find the three axes that maximally separate left vs. right correct trials between movement onset and reward (axis 1, shown in Figure 4g), movement onset time vs. the time of reward in all correct trials (axis 2), and the time of motor preparation vs. motor execution (trial start vs. mid-movement, axis 3). We orthonormalized the three axes and projected raw and reconstructed data onto the new, three-dimensional basis Figure 4h.

We then measured the distance ratio between trials of the same vs. between trials of a distinct trial class (left vs. right) in the full neural space. For the reconstructed vs. the full data set, we averaged neural activity over a 650 ms window centered at movement onset and measured the Euclidean distance of the population response in each trial to the trial-averaged population response in its own trial type, compared to the Euclidean distance to the average population response of the respective other trial type: Δ_between_/Δ_within_, where 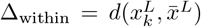 is the Euclidean distance between population vectors in each left trial to the mean population vector across all left trials (and vice versa for right trials), and 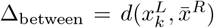 is the Euclidean distance of population vectors in each left trial to the mean population vector across all right trials (and vice versa for right trials).

### 4.7 Dataset 3: Electrophysiology across many brain regions during perceptual decision making

#### 4.7.1 Description of the dataset

The third analyzed dataset comprised recently published multi-region Neuropixel recordings (*N* = 303) in a mouse performing a perceptual decision making task [IBL et al., 2021]. In the task, mice were presented a grating patch image with varying contrast (0%, 25%, 35%, 50% or 100%), shown on the left or right sides of a screen. The mice were trained to move the image to the center of the screen using a steering wheel within a 60 s period in order to receive a sugary water reward. A correct response was registered if the stimulus was moved to the center, and an incorrect response if the stimulus was moved to the border of the screen. We selected a single example mouse (subject CSHL049 from the openly released ephys data repository).

#### 4.7.2 Additional preprocessing

We binned single-neuron spiking events in 10 ms windows. Due to the variable response times across trials, we piecewise linearly warped data between stimulus onset and reward delivery or respectively, timeout onset, to correspond to the median interval length, and clipped the trial period to start 1 s before stimulus onset and to end 2 s after reward delivery or timeout onset. We smoothed data with a Gaussian filter with *σ* = 20ms and rescaled the activity of each neuron to a minimal and maximal value of 0 and 1 over all trials. We excluded neurons with mean firing rates below 0.2 Hz, leading to a total of *N* = 221 neurons analyzed out of *N* = 303 neurons recorded. Brain regions included visual cortex (VIS: anterior layers 2/3, 4, 5, 6a and 6b as well as anteromedial layers 2/3, 4, 5, and 6a; *N* = 85 neurons), hippocampal regions CA1 (*N* = 32 neurons) and dentate gyrus (DG: molecular, polymorph, and granule cell layers; *N* = 21 neurons), thalamus (TH, including posterior limiting nucleus and lateral posterior nucleus; *N* = 18 neurons) and the anterior pretectal and midbrain reticular nucleus (APN, *N* = 22 neurons, and MRN, *N* = 35 neurons) of the midbrain. In total, the resulting data tensor had dimensions *N* = 221, *T* = 350, and *K* = 831.

#### 4.7.3 Visualization of sliceTCA weights

In Figure 5b, we scaled the rows of the neuron-time slices to a [0,1] interval to highlight differences in the timing of peak activity between neurons. We then reordered neuron-time slices by peak activity within each region for each slice type separately, to show characteristic differences between neural correlates of behavioral variables. Trial-time slices were regrouped by trial type to show region-specific representations of task variables. Finally, neuron-trial slices were reordered by average weights across the first 100 trials for each neuron within a region.

#### 4.7.4 Reconstruction performance and component weights

For each neuron, we estimated the goodness of fit of the sliceTCA reconstruction as:

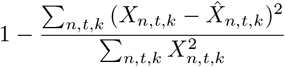

We then quantified the contribution of the neuron-slicing components on the total sliceTCA reconstruction for each neuron n as the following ratio:

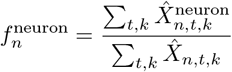

where 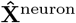 describes the partial reconstruction of the data tensor from only the neuron-slicing components. We similarly defined the contributions of the time- and trial-slicing components to the sliceTCA reconstruction of each neuron *n* as 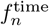 and 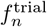.

### 4.8 Code availability

A GPU accelerated Python library for the sliceTCA data analysis pipeline (including preprocessing, model selection, model optimization, and visualization of components) is available at https://github.com/arthurpe/slicetca. In addition, the code necessary for reproducing main analyses will be published in a separate Github repository upon publication.

## 4.9 Acknowledgements

We thank Joao Barbosa, Matthias Hennig, Kishore Kuchibhotla, Ashok Litwin-Kumar, Arno Onken, Yann Sweeney, and the members of the Cayco Gajic lab for comments on the manuscript. We additionally thank Angus Chadwick for helpful discussions on an early stage of the manuscript, and Mark Wagner for sharing his data. We are also grateful to the IBL, the Churchland/Shenoy labs, and the Neural Latents Benchmark project for making their processed and curated data freely available. This work was funded by EMBO (H.S., ALTF 471-2021) and the Agence Nationale de la Recherche (N.A.C.G., ANR-20-CE37-0004; ANR-17-EURE-0017).

## 4.10 Author contributions

A.P., H.S., and N.A.C.G. conceptualized the project. A.P. and H.S. designed the data analysis pipeline. A.P. and H.S. performed data analysis investigations. A.P., H.S., and N.A.C.G designed the feedforward model. A.P. wrote the mathematical notes. N.A.C.G. wrote an initial draft of the manuscript, which all authors reviewed and revised. N.A.C.G. supervised the project.

---

**Supplementary Figure 1:**
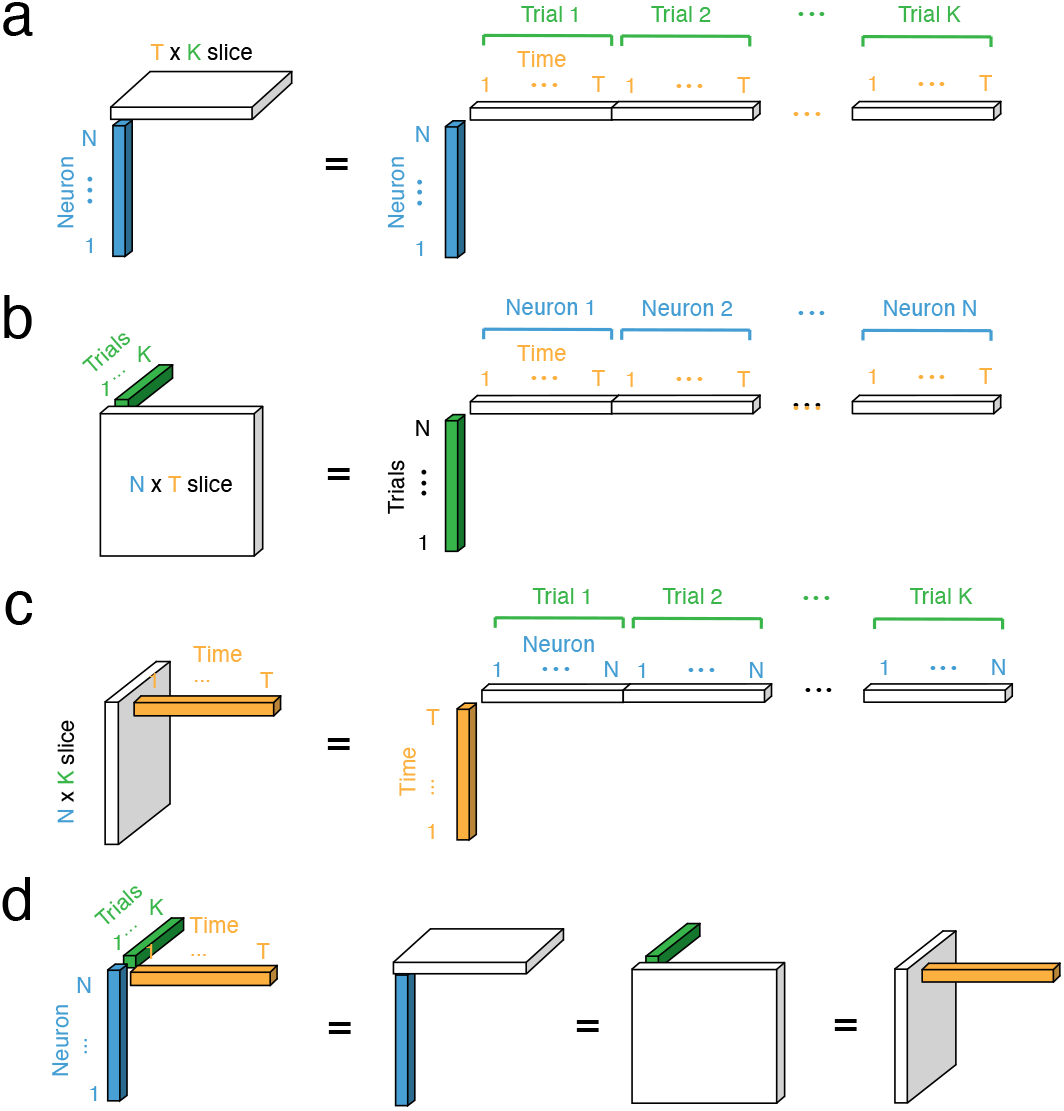
Tensor decomposition, matrix factorizations, and covariability classes. **a.** A neuron-slicing component can be converted into a rank-1 matrix by unfolding the slice into a row vector (or equivalently, by unfolding the reconstructed 3-tensor into a matrix). A sliceTCA model comprised only of neuronslicing components (i.e., *R*_time_ = *R*_trial_ = 0) is equivalent to applying matrix factorization after unfolding the data tensor into an *N* × *KT* matrix (e.g., as in ‘trial-concatenated PCA’). Therefore, a neuron-slicing component can be interpreted as latent dynamics that varies over trials but which is shared across neurons. **b-c.** The trial-slicing (**b.**) and time-slicing (**c.**) components can similarly be converted to rank-1 matrices. These components represent shared variability across trials and time, respectively. **d.** A TCA component can be represented as any of the three sliceTCA components by defining the slice be the outer product of two of the loading vectors. This means that a single TCA component simultaneously corresponds to shared variability across neurons, time, and trials.

**Supplementary Figure 2:**
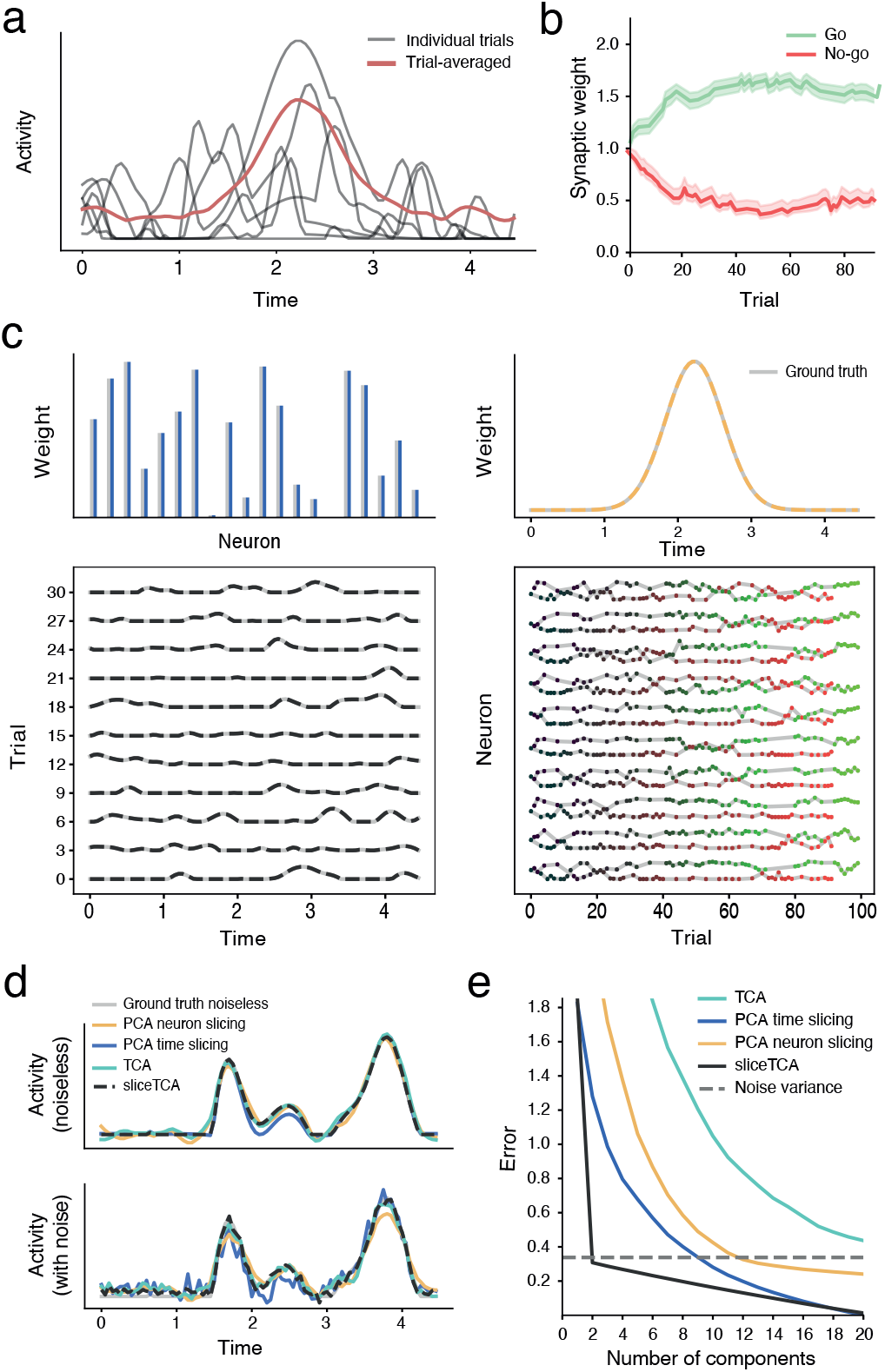
Feedforward model of perceptual learning. **a.** Sample of the activity of a single neuron over six example trials (grey) as well as trial-averaged activity (red). **b.** Evolution of the mean Go and No-go weight over learning. Shading represents the standard error of the mean. **c.** Recovered neuron-slicing (left) and timeslicing (right) components, plotted alongside ground truth values (grey). Weights for each neuron in the slice of the time-slicing component are plotted separately for Go (green) and No-go (red) trials. The sliceTCA decomposition captures the ground truth exactly. **d.** Single neuron activity is better captured by sliceTCA with (bottom) and without (top) white noise added to the data tensor. **e.** Mean squared error of decomposing the activity of the feedforward model with added white noise for sliceTCA, TCA, and PCA as a function of the number of components used in the model. The dashed line represents the mean squared deviation between the noisy and the noiseless model. Adding white noise does not affect the performance of sliceTCA relative to TCA or PCA.

**Supplementary Figure 3:**
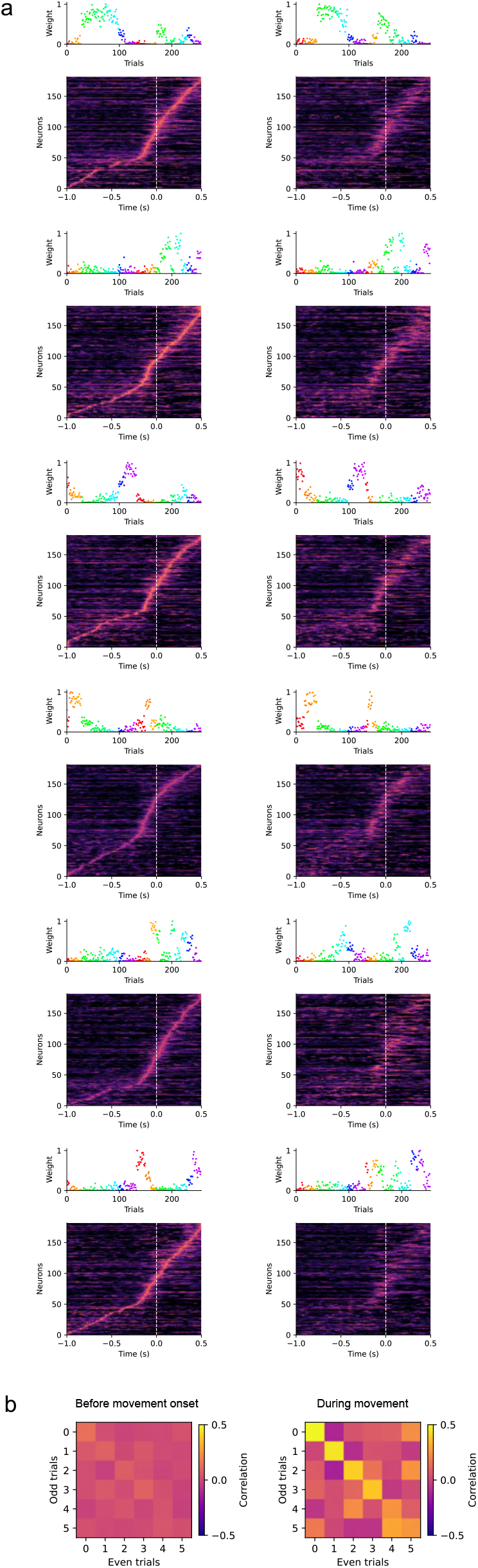
Cross-validation of neuron-specific sequences identified in trial-slicing components. **a** Components identified by trial-slicing NMF (*R*_neuron_ = 6) applied to the motor cortical reaching dataset (Figure 2). To validate the sequences we separated the data into even (left) and odd (right) trials and fit the model separately on each of the two datasets (only 6 rather than 12 components to avoid overfitting since we halved the datasets). Components from the two models were matched by hand based on similarity of the trial loading vector. In each row of the panel, neurons in the slices are sorted identically (according to the latency of peak activity in even trials). Neural sequences during movement (dashed white line indicates movement onset) are reproduced in the two independent groups of trials, while pre-motor sequences are not matched. **b** To quantify how reliably movement-related and pre-motor sequences could be identified across splits,for each component we calculated the Pearson correlation for each pair of slices across the two models, separately for pre-movement onset (−1 to −0.2 s) and during movement (−0.1 −0.5 s) periods.

**Supplementary Figure 4:**
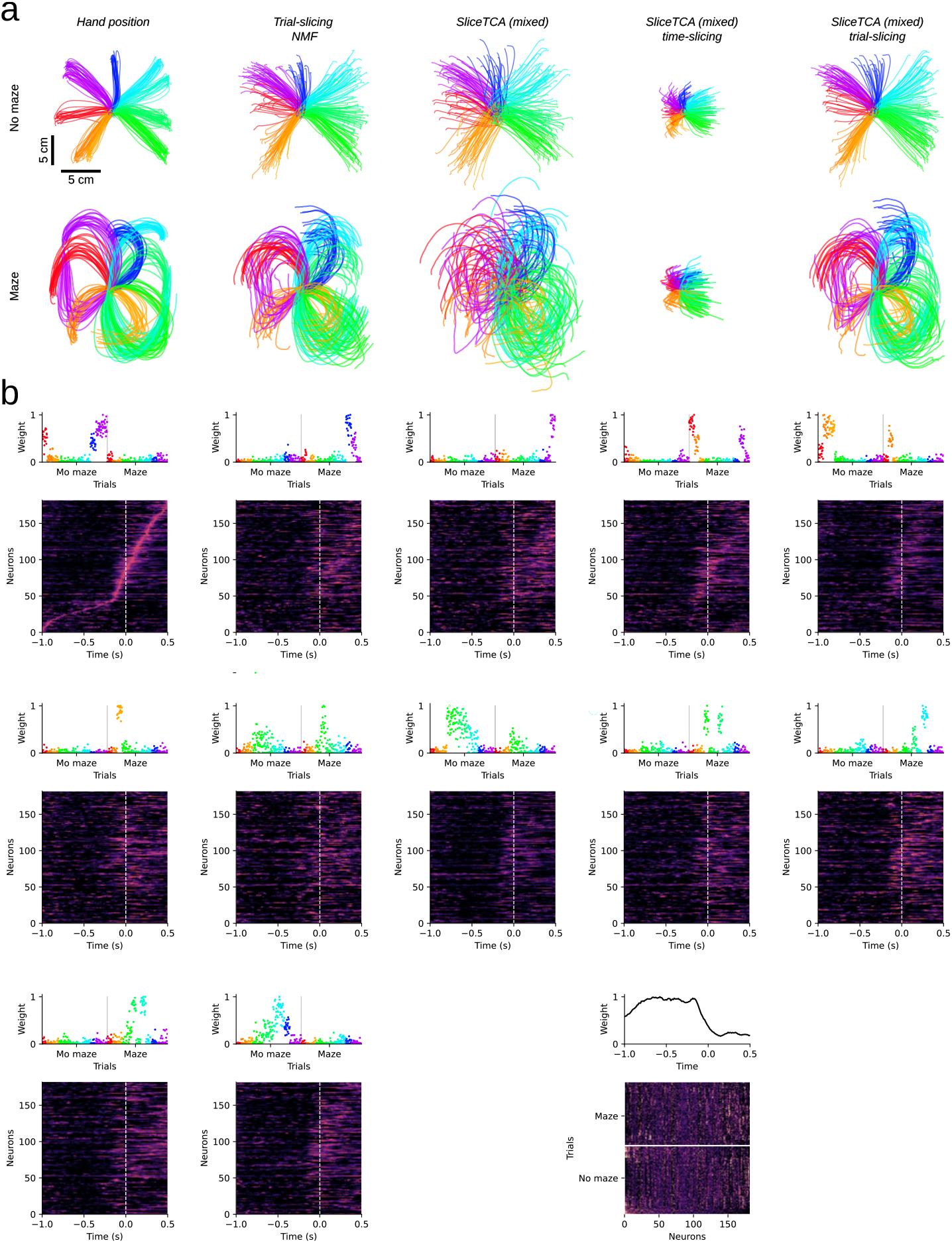
Analysis of mixed covariability model in decoding analysis of reaching kinematics. **a.** Ordinary least square decoding of the hand velocity from different decompositions. For comparison, hand position and the mapping from the reconstruction of the the trial-slicing only NMF model (*R*_neuron_ = 0, *R*_time_ = 0, *R*_trial_ = 12) are reproduced from Figure 2. Also shown are the decoding performances using the fit of the mixed covariability model (*R*_neuron_ = 0, *R*_time_ = 1, *R*_trial_ = 12), based on the full reconstruction of all components in the model, as well as the partial reconstruction of the data tensor from only the time-slicing component or only the trial-slicing components. Note that adding the single time component to the mixed model before decoding decreases performance, indicating that it captures variability in the neural data that is not directly relevant to movement. **b.** The 12 trial-slicing components of the mixed covariability model. In the slices, neurons are ordered according to latency of peak activation in the first component. These components seem to capture similar reach directions as seen by the loading vector weights having a bell-shaped tuning curve to certain preferred angles. Furthermore, some components seem to capture specifically maze-related neural variability. **c.** The time-slicing component of the mixed covariability model, reproduced from Figure 2d for reference.

**Supplementary Figure 5:**
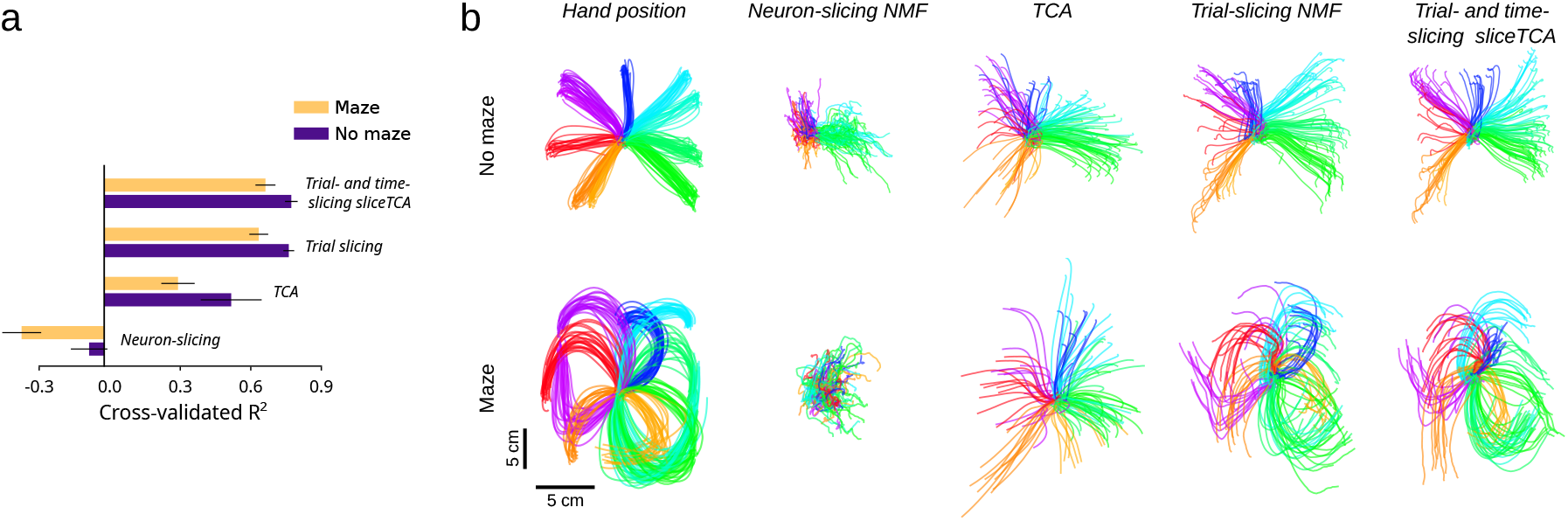
Simultaneous dimensionality reduction and decoding cross-validation in the motor cortex reaching dataset. **a.** We also performed the decoding analysis with a stricter form of cross-validation (cf. Figure 2e). We first split the data in train and test sets with equal number of trials (randomly chosen). We fit the respective dimensionality reduction method to both sets separately, and trained the decoder only on the training dataset. The *R*^2^ reported is that of the decoded hand movement on the test dataset. The operation is repeated by swapping which half is used for fitting and testing (2-fold validation), and repeated over 10 permutations. Error bars indicate standard deviation. Note that in Figure 2, to allow comparison with other methods, we applied the respective dimensionality reduction on the full dataset, before splitting into train- and test-sets for the decoding analysis. Moreover, to avoid overfitting the decoder to the smaller training dataset (compared to Figure 2a), we here use ridge regression for decoding instead of OLS (*L*_2_ regularization coefficient equal to 5 as determined by hyperparameter tuning). **b.** Decoded hand movement on an example test set.

**Supplementary Figure 6:**
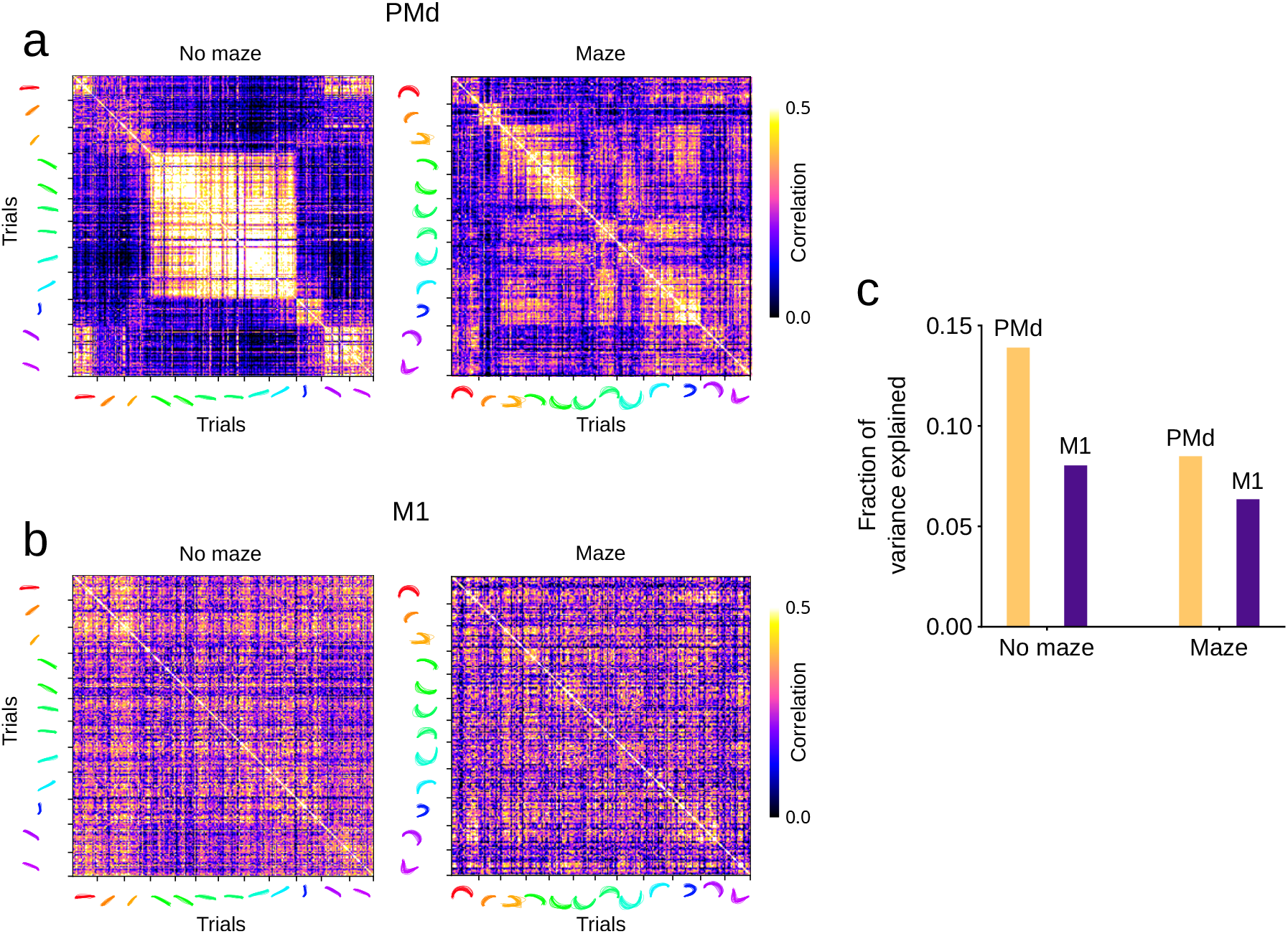
Reach condition-specific structure in motor preparatory information is observed in time-slicing weights of PMd, but not of M1. **a.** Correlation matrix between neural encoding weights for pairs of trials in the time-slicing matrix, showing similarity of PMd neuron weights for trials with similar reach direction and curvature. Reproduced from Figure 2f. **b.** In contrast, M1 neuron weights reveal a lack of structure in its correlation matrices. **c.** Average fraction of the variance of PMd and M1 neuron activity explained by single time-slicing component, separated by maze and no maze conditions.

**Supplementary Figure 7:**
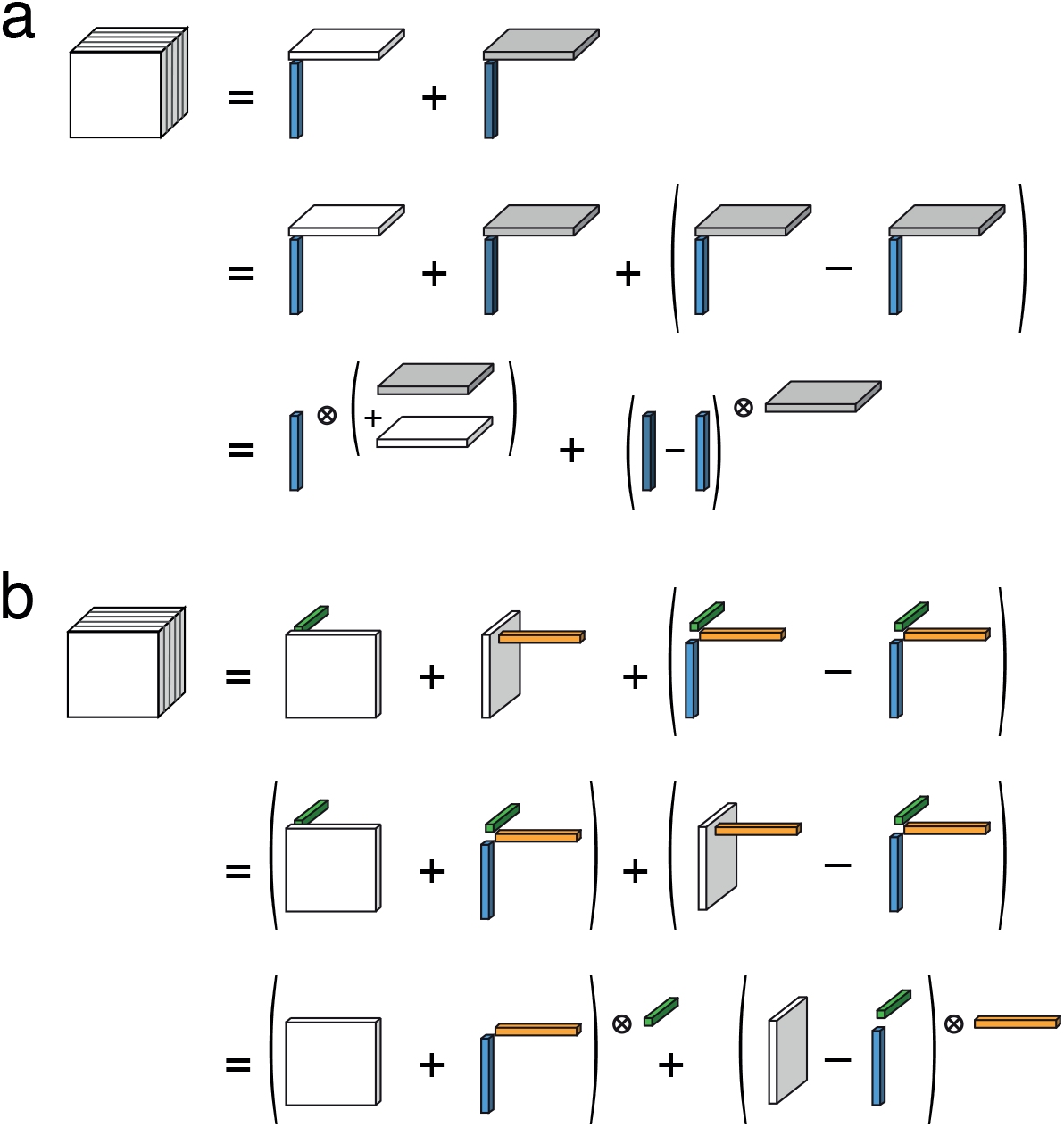
Schematic of the two sliceTCA invariance classes. **a.** Example of a within-slicetype invariant transformation in a slice-rank-2 tensor formed by adding and subtracting a slice-rank-1 component with the same loading vector as component 1 and the same slice as component 2. These terms can be absorbed into the original two components, resulting in two equivalent decompositions. **b.** Example of a between-slice-type invariant transformation in a slice-rank-2 tensor. Here, a rank-1 tensor constructed by the two components’ loading vectors (green and yellow) and a third free vector (blue) is added and subtracted. These terms can be absorbed into the original two components, resulting in two equivalent decompositions.

**Supplementary Figure 8:**
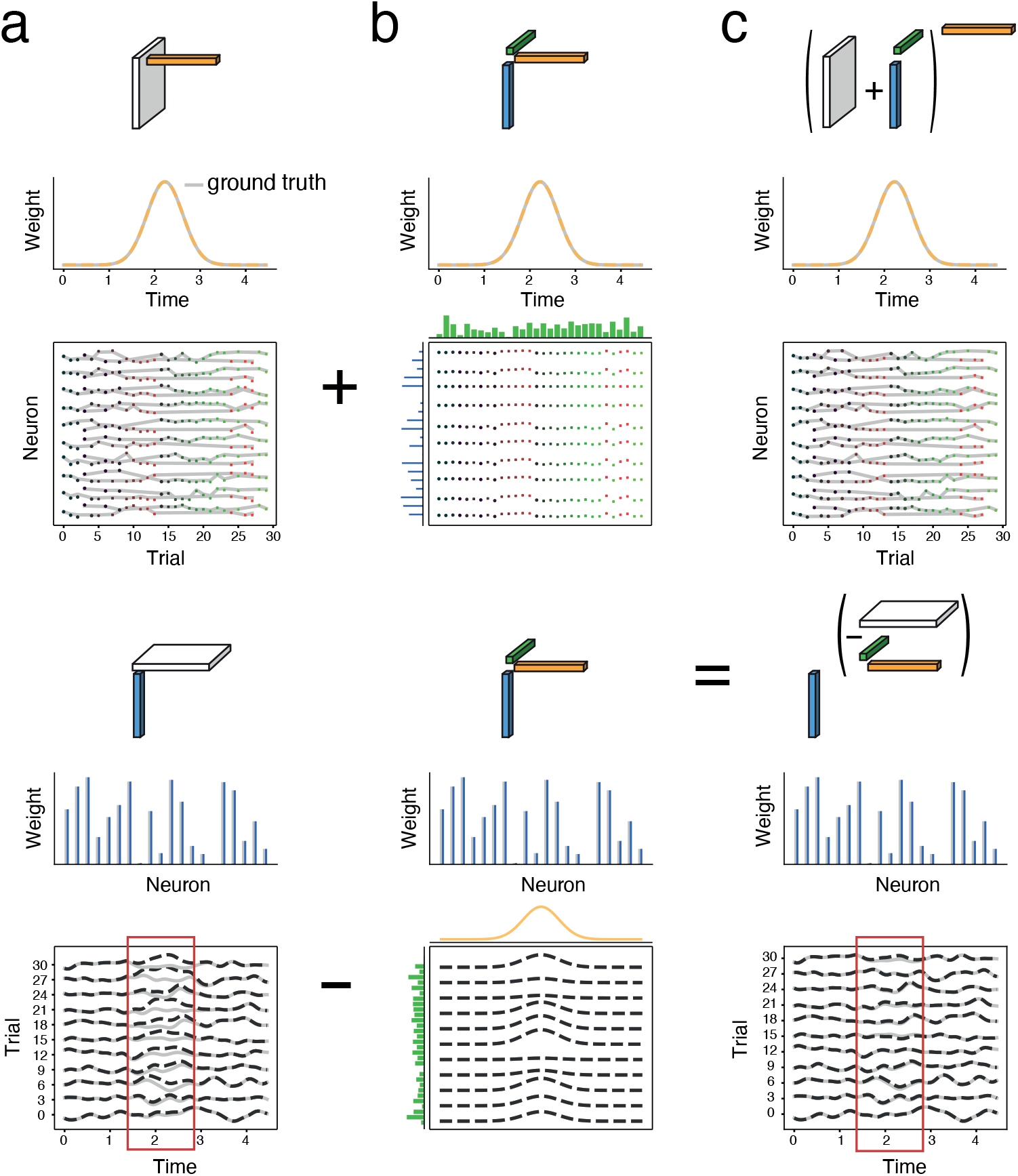
Between-slice-type invariance in the feedforward model. **a.** The activity of the feedforward model (Supplementary Figure 2) can be decomposed into the sum of a time-slicing component (top) and a neuron-slicing component (bottom). In this example, the Gaussian Process is not rectified and the data is not non-negative, leading to bleedthrough between the two components due to this invariance class (bottom, red rectangle shows discrepancy between model and ground truth). This invariance class was not observed in the original non-negative model (Figure 1h-j, Supplementary Figure 2) because there are fewer permissible transformations when the factors are more constrained. **b.** Example of a rank-1 tensor that can be passed between the two components. Two of the loading vectors are identical to the loading vectors of the componengs in **a**. The third is unconstrained. **c.** By choosing the trial loading that minimizes a specified objevtive function, a unique solution can be found. Here the correlation between the activity of trial-neuron pairs was minimized in the neuron slicing component, resulting in a fit that matches the ground-truth values.

**Supplementary Figure 9:**
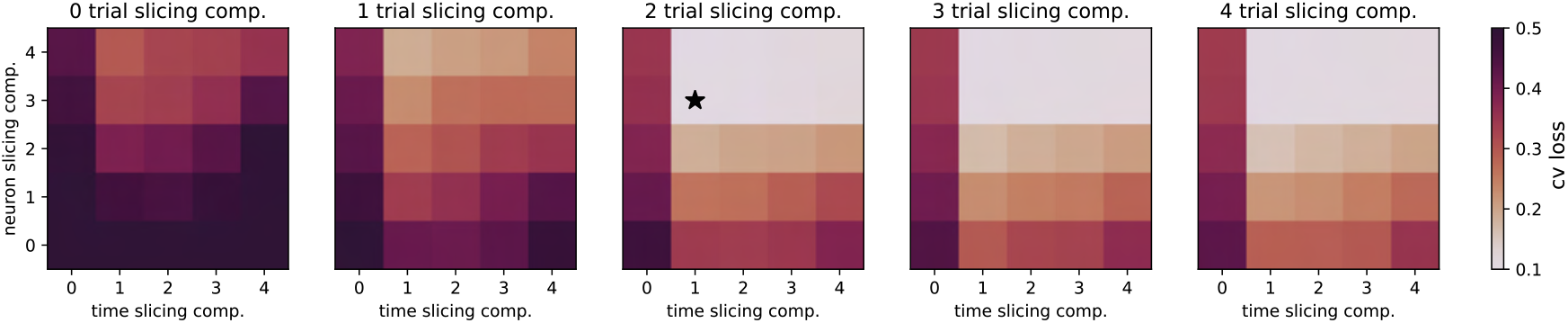
Validation of cross-validated model selection procedure. For ten random seeds, data tensors were split in train- and test-data (80% and 20%, respectively), following a blocked masking procedure as described in Methods. Grid shows mean squared error loss for different numbers of components of each slice type, averaged across ten cross-validation folds. The ground truth model was a fit to the cortico-cerebellar data with decomposition of *R*_neuron_ = 3, *R*_time_ = 1, *R*_trial_ = 2 with independent Gaussian noise (*σ* = 0.1) added to each tensor entry. The gridsearch recovers the correct number of components of each slice type (black star indicates optimal model).

**Supplementary Figure 10:**
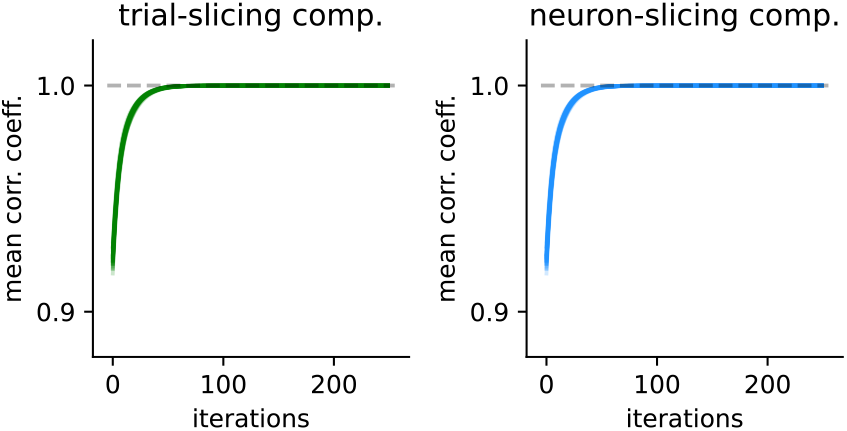
Validation of hierarchical model optimization procedure. Results of running the tensor-passing optimization step (L2 in Figure 3c) on the decomposition shown in Figure 4, with 10 random seeds, so that the free vector of the rank-1 tensor was initialized differently each time. During optimization, the free vector is optimized to minimize the L2-norm of slices. Over 250 iterations, the optimization procedure lead to a unique solution, observable in a perfect correlation between slice weights both for trial- (green, left) and neuron-slicing components (blue, right). Each transparent line is the pairwise correlation between two solutions found based on 10 different initialization seeds.

**Supplementary Figure 11:**
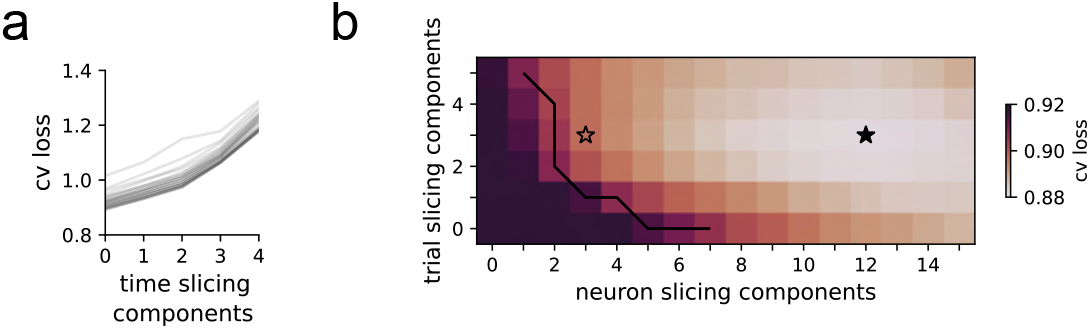
Cross-validated model selection for cerebello-cortical imaging dataset. **a.** The cross-validated loss as a function of the number of time-slicing components. Transparent grey lines show different combinations of 0-5 neuron- and 0-5 trial-slicing components. This indicates that adding any time-slicing components leads to an overfitting of the data. **b.** The cross-validated loss as a function of the number of neuron- and trial-slicing components (for 0 time-slicing components). The filled star indicates the minimum loss (i.e., the optimal model), while the black line indicates the 80% loss elbow. The hollow star indicates the model selected for further analysis in Figure 4.

**Supplementary Figure 12:**
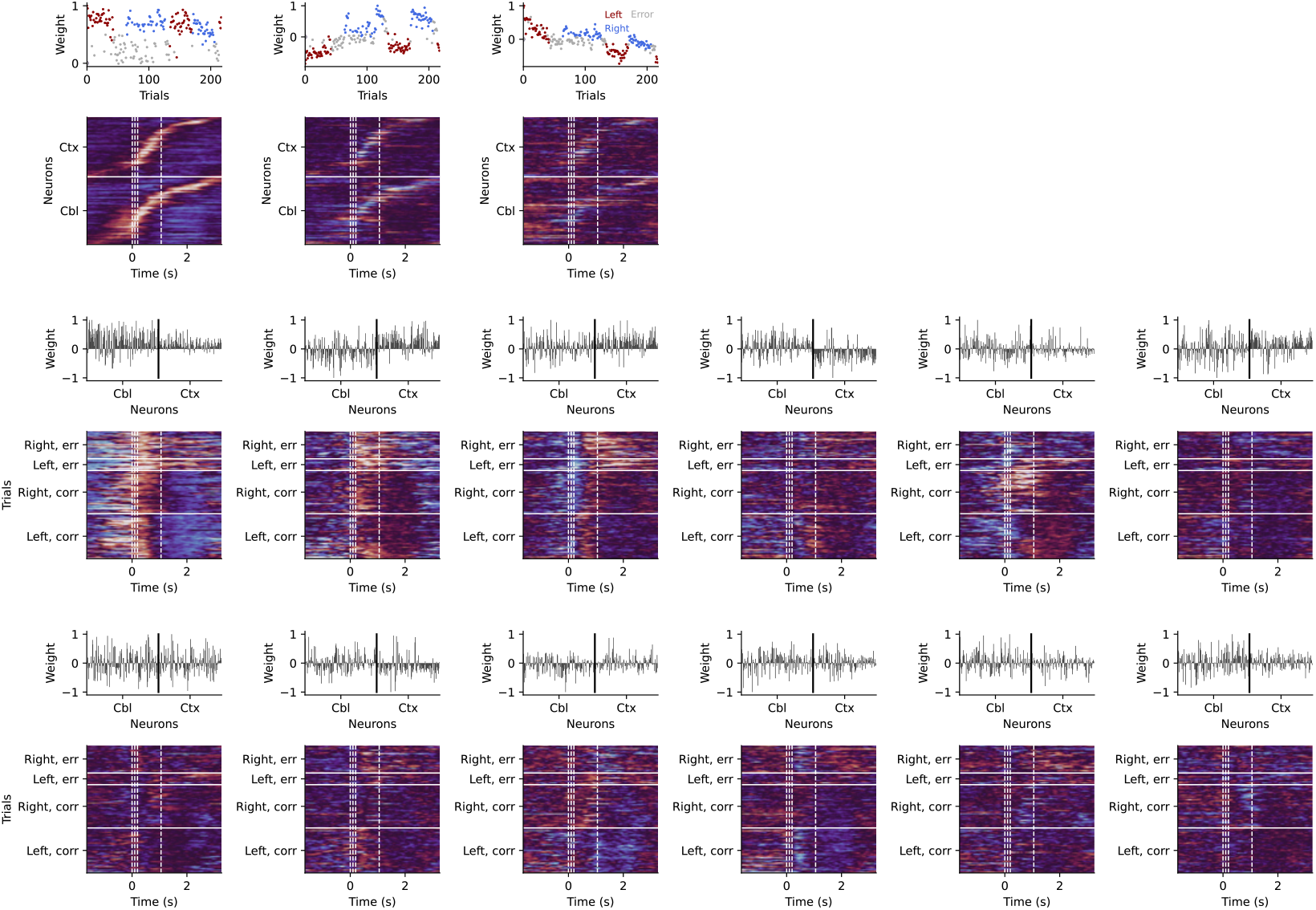
Optimal model of cerebello-cortical imaging dataset. The three trial-slicing and twelve neuron-slicing components of the optimal model found from the cross-validated grid search (Supplementary Figure 11). Note that the first three neuron-slicing components, as well as all trial-slicing components, are highly similar to the components in the selected model in Figure 4b,c.

**Supplementary Figure 13:**
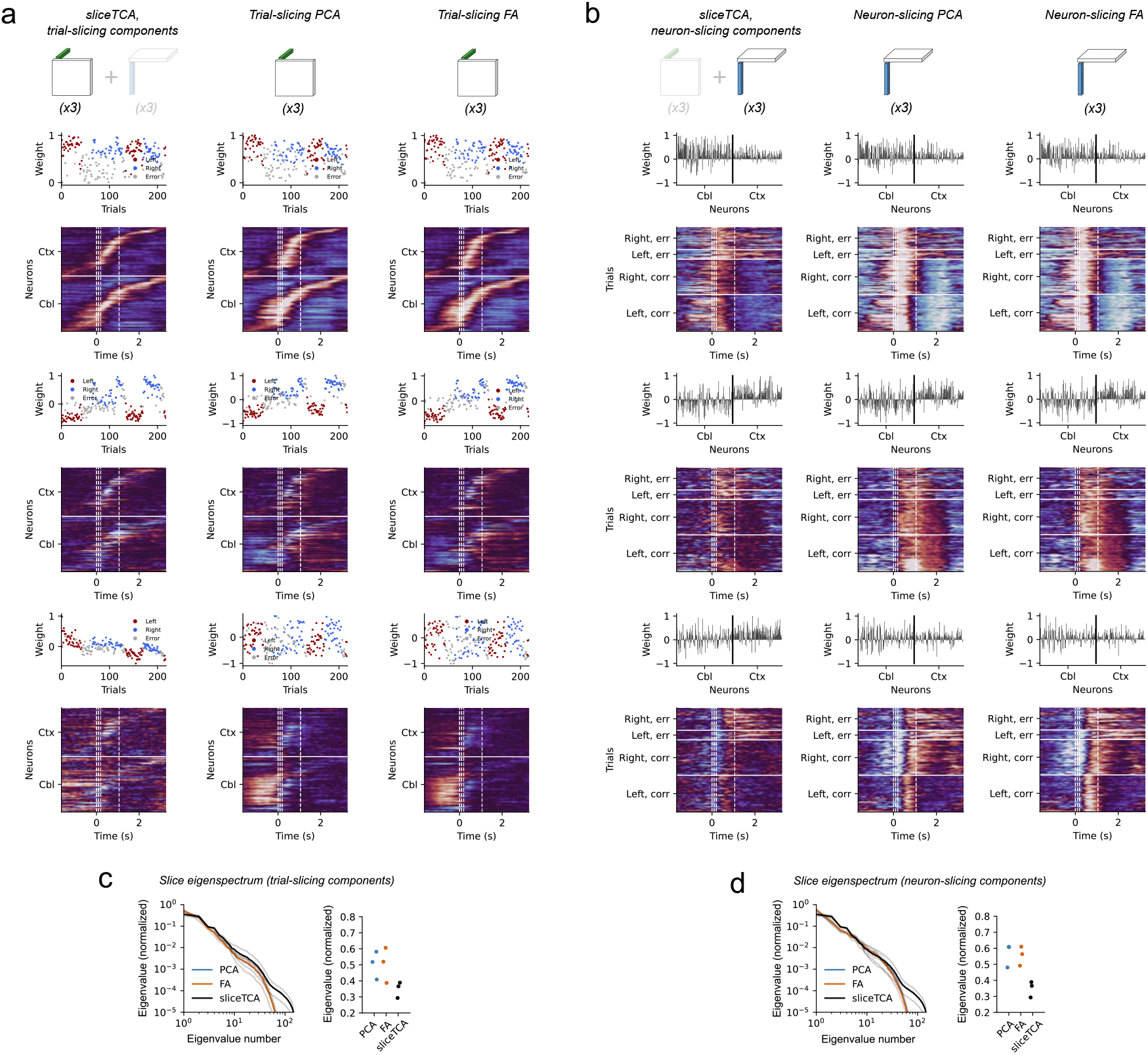
PCA and FA components in neuron- and trial-slicing components. **a.** The three trial-slicing components of the sliceTCA model selected in Figure 4b,c (left), plotted alongside the first three trial-slicing PCA components (middle), and three trial-slicing FA components (right). All slices are ordered by peak neuron activity. Note that the sliceTCA slices appear to be higher rank (as seen by the fact that they appear more diagonally weighted; Figure 4e)). SliceTCA also seems to better capture adaptation-like dynamics in the third component, and better cluster trial types compared to PCA and FA. **b.** Neuron-slicing component of the same sliceTCA decomposition (left), plotted alongside the first three neuron-slicing PCA components (middle), and three neuron-slicing FA components (right). All slices are ordered by trial condition-outcome pairs. Again, the sliceTCA slices appear to be higher rank. **c.** Eigenspectrum analysis for slices of trial-slicing components. Same as Figure 4e. **d.** Eigenspectrum analysis for slices of neuron-slicing components.

**Supplementary Figure 14:**
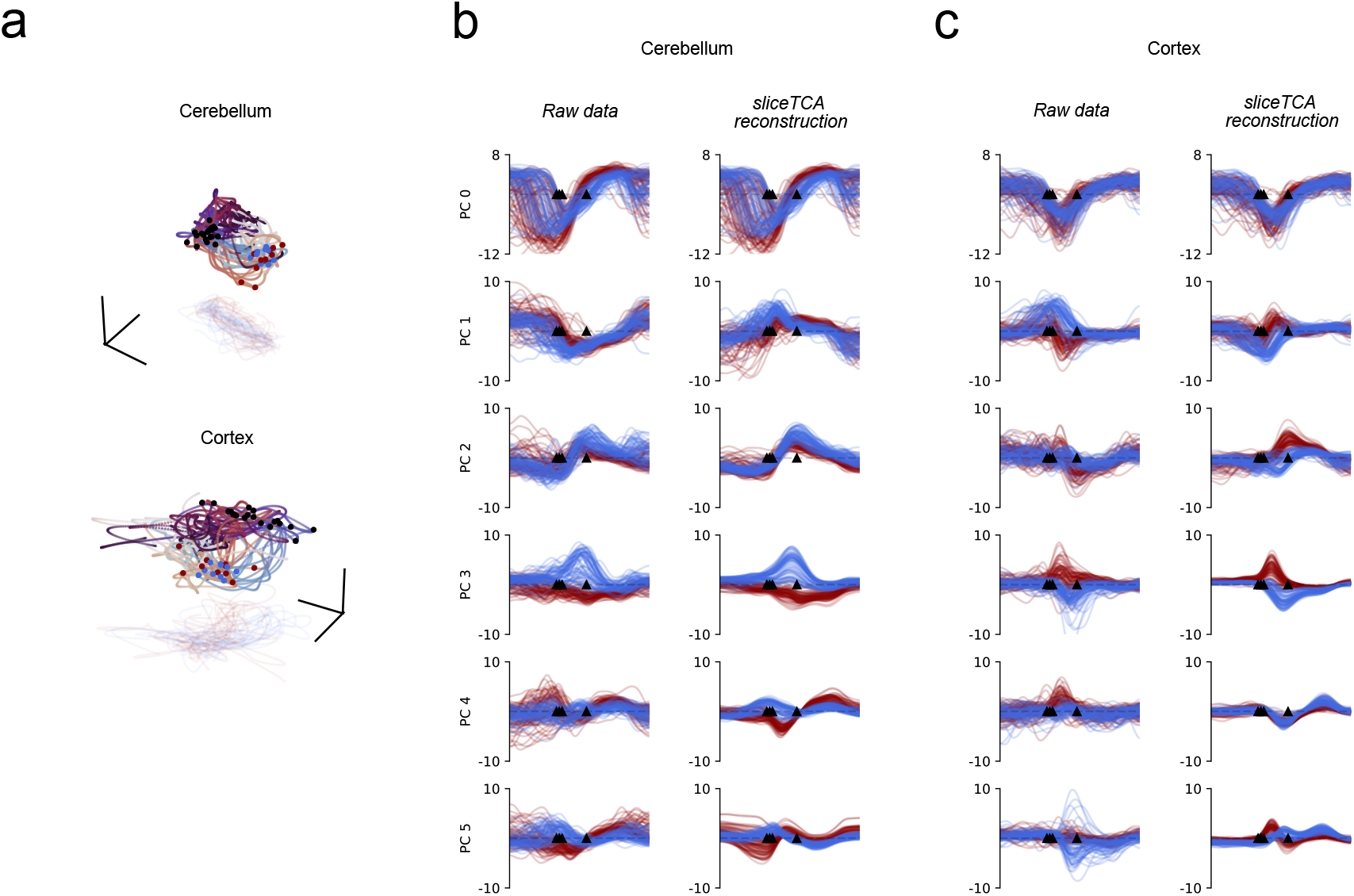
Raw data manifolds projected onto LDA axes and principal components in raw data and sliceTCA reconstructions. **a.** Projection of raw data onto LDA axes that separate left from right trials between movement onset and reward; pre-motor and mid-movement; or movement onset vs. reward time, respectively (cf, Figure 4h; here, LDA axes are found from raw data) **b.** Projection of raw data (left) and sliceTCA reconstructions (right) of cerebellar population activity onto the first six principal components found on raw data (left) and sliceTCA reconstructions (right) for all correct trials. Higher principal components found in sliceTCA reconstructions (from third PC onwards) appear smoother than those found in raw data, and separate trials of different reach direction more clearly. **c.** Same as b for the cortical population.

**Supplementary Figure 15:**
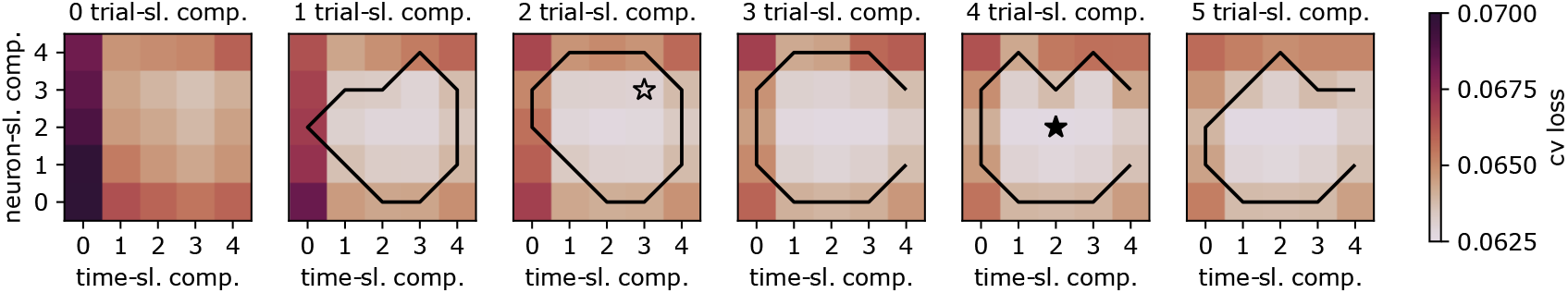
Cross-validated model selection for multi-region Neuropixel dataset. For 10 random seeds, data tensors were split in train- and test-data (80% and 20%, respectively), following a blocked masking procedure as described in Methods. Full black star marks the optimal model, and hollow black star marks the selected model analyzed in Figure 5. Black thresholds describe a 95 % loss elbow.

**Supplementary Figure 16:**
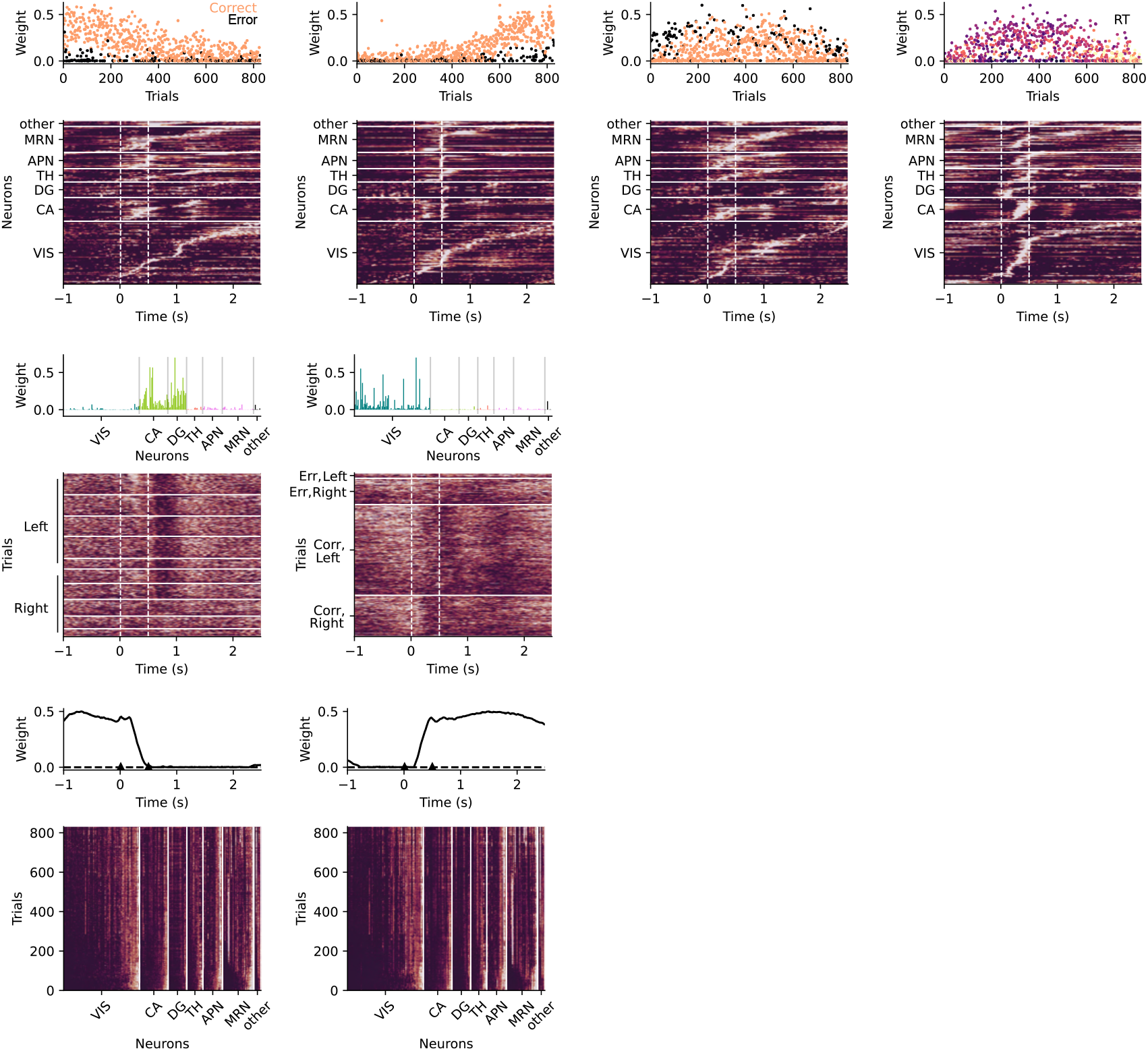
Optimal model of multi-region Neuropixel dataset. Components of the optimal model identified by the cross-validated grid search (Supplementary Figure 15), with 4 trial-slicing components, 2 neuron-slicing and 2 time-slicing components. Similarly to decomposiiton selected for Figure 5, this model identified time-slicing components related to correct vs. error trials, and a RT-related component. Neural activity related to correct vs. error encoding differed between late and early trials. Moreover, the optimal model grouped CA and DG together in a neuron-slicing component, and lacked a distinction between reward vs. post-reward related time courses in the two time-slicing components.

**Supplementary Figure 17:**
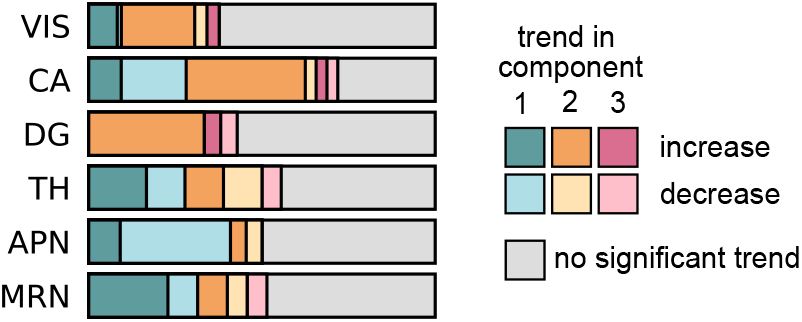
Time-slicing components do not sum to a flat baseline. To determine whether the three time-slicing components found by sliceTCA reconstructed an increasing baseline for most neurons or whether responses in single task periods changed more strongly than others, we compared the slope of linear trends over trials (for each neuron separately) across the three time-slicing components. Specifically, we fitted linear models with a multiplicative term for each neuron: *w_k,r_* = *β*_0_ + *β*_1_*k* + *β*_2_*δ*_*r*=2_ + *β*_3_*δ*_*r*=3_ + *β*_4_*kδ*_*r*=2_ + *β*_5_*kδ*_*r*=3_ + *ε_k,r_* where *w_k,r_* is the weight in trial *k* for component *r*, and *δ*_*r*=2_ = 1 if the weight belongs to component 2, *δ*_*r*=2_ = 0 otherwise (same for *δ*_*r*=3_, whereas component 1 is the reference class). We performed analyses of variance for each neuron to test whether multiplicative terms explained a significant part of variance (at a Bonferroni-corrected significance level a = 0.05 with *N* =221 neurons). We discarded neurons with non-significant multiplicative terms (grey). For neurons with significant differences in the rate of change across time-slicing, we compared rates of change between components. Each neuron was classified by the highest absolute change coefficient and its sign, leading to six classes for three time-slicing components.

**Supplementary Figure 18:**
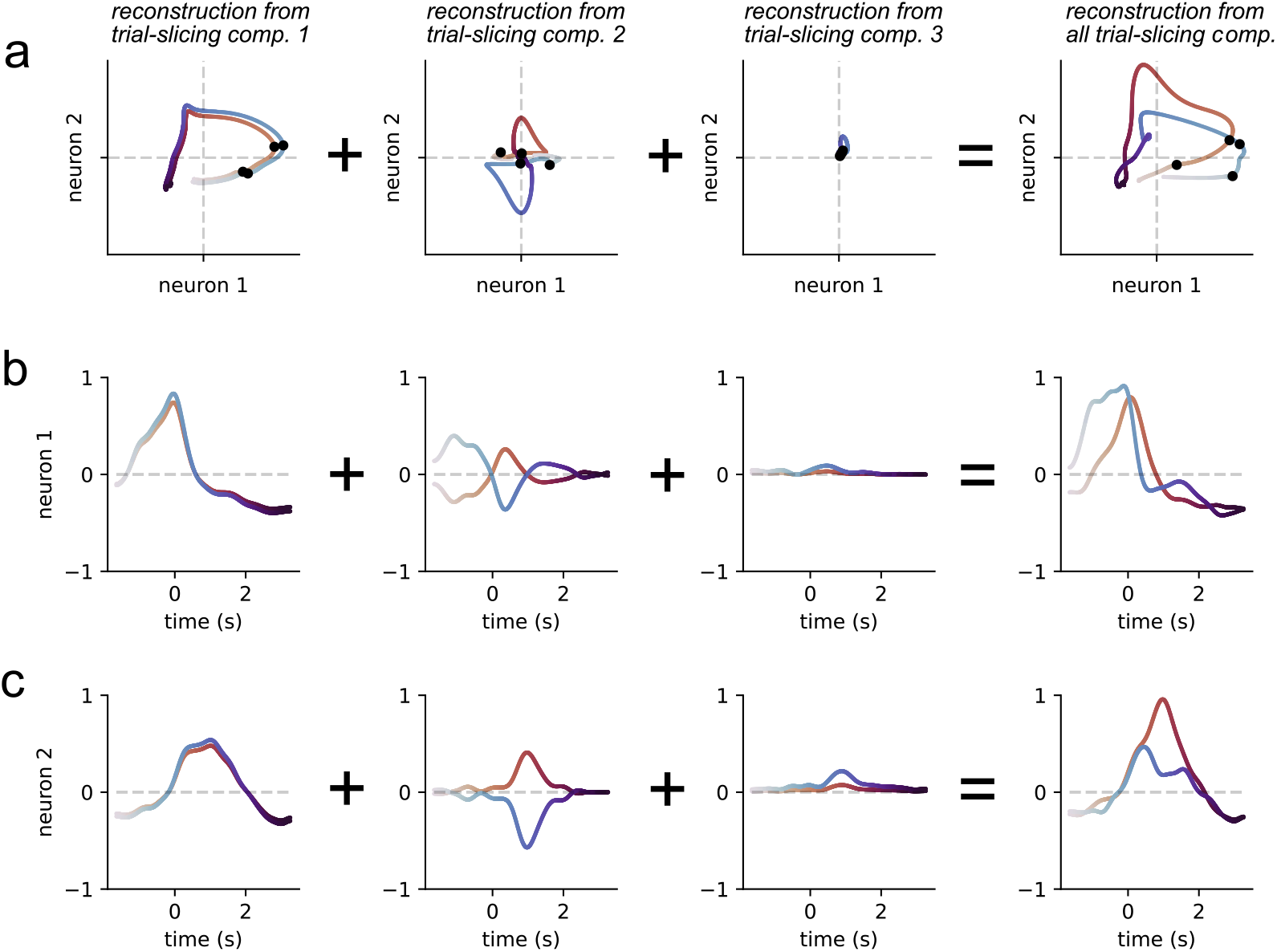
Example of reconstruction of multiple trial-slicing components. **a.** Reconstruction of neural trajectories from three trial-slicing components of the dataset presented in Figure 4. For single components, neural trajectories in the two trials are scaled versions of another. However, the full trial-slicing reconstructions (right) are not simply scaled versions of each other since the three components have different scaling factors. Therefore multiple trial-slicing components can lead to more complex latent dynamics than shown in Figure 6. **b-c.** Single neuron reconstructions plotted against time for each component and the full trial-slicing reconstruction.

**Supplementary Figure 19:**
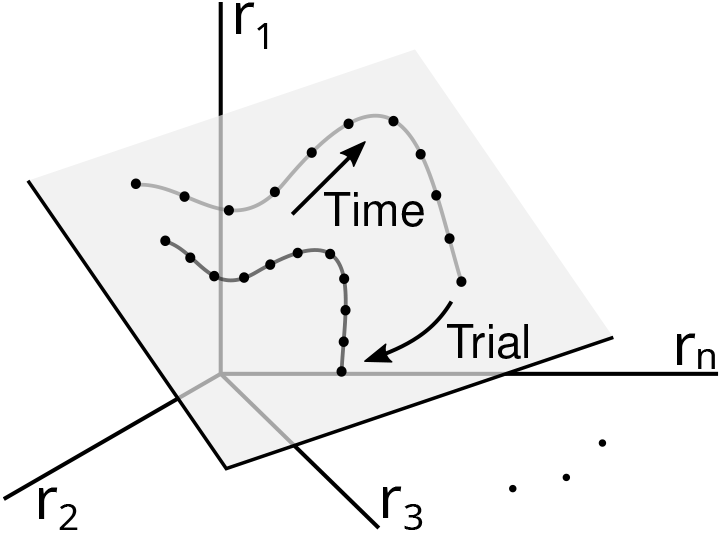
Geometric interpretation of TCA components in neural activity space. Similarly to sliceTCA and matrix factorization methods (Figure 6), TCA can be interpreted as uncovering latent variables embedded in neural activity space. Since a TCA component lies at the intersection of the three covariability classes (Figure 1g), it must obey all three constraints described in Figure 6. In other words, the latent dynamics must be embedded in a fixed R-dimensional subspace and can only change in amplitude (not the shape of the dynamics) along the corresponding latent variable across trials.

## References

M. B. Ahrens, J. M. Li, M. B. Orger, D. N. Robson, A. F. Schier, F. Engert, and R. Portugues. Brain-wide neuronal dynamics during motor adaptation in zebrafish. Nature, 485(7399):471–477, 2012.

R. Amo, S. Matias, A. Yamanaka, K. F. Tanaka, N. Uchida, and M. Watabe-Uchida A gradual temporal shift of dopamine responses mirrors the progression of temporal difference error in machine learning. Nature Neuroscience, pages 1–11, 2022.

M. Balasubramanian and E. L. Schwartz. The isomap algorithm and topological stability. Science, 295 (5552):7–7, 2002.

E. Balzani, J. P. Noel, P. Herrero-Vidal, D. E. Angelaki, and C. Savin. A probabilistic framework for task-aligned intra-and inter-area neural manifold estimation, 2022. URL https://arxiv.org/abs/2209.02816.

M. Belkin and P. Niyogi. Laplacian eigenmaps for dimensionality reduction and data representation. Neural computation, 15(6):1373–1396, 2003.

M. Bläser, C. Ikenmeyer, V. Lysikov, A. Pandey, and F. Schreyer. Variety membership testing, algebraic natural proofs, and geometric complexity theory. CoRR, abs/1911.02534, 2019. URL http://arxiv.org/abs/1911.02534

J. D. Carroll and J.-J. Chang. Analysis of individual differences in multidimensional scaling via an n-way generalization of “eckart-young” decomposition. Psychometrika, 35(3):283–319, 1970.

R. Chaudhuri, B. Gerçek, B. Pandey, A. Peyrache, and I. Fiete. The intrinsic attractor manifold and population dynamics of a canonical cognitive circuit across waking and sleep. Nature neuroscience, 22(9): 1512–1520, 2019.

M. M. Churchland, J. P. Cunningham, M. T. Kaufman, J. D. Foster, P. Nuyujukian, S. I. Ryu, and K. V. Shenoy. Neural population dynamics during reaching. Nature, 487:51–56, 2012.

J. P. Cunningham and B. M. Yu. Dimensionality reduction for large-scale neural recordings. Nature neuroscience, 17(11):1500–1509, 2014.

L. N. Driscoll, N. L. Pettit, M. Minderer, S. N. Chettih, and C. D. Harvey. Dynamic reorganization of neuronal activity patterns in parietal cortex. Cell, 170(5):986–999, 2017.

E. L. Dyer, M. Gheshlaghi Azar, M. G. Perich, H. L. Fernandes, S. Naufel, L. E. Miller, and K. P. Körding. A cryptography-based approach for movement decoding. Nature biomedical engineering, 1(12):967–976, 2017.

S. Ebrahimi, J. Lecoq, O. Rumyantsev, T. Tasci, Y. Zhang, C. Irimia, J. Li, S. Ganguli, and M.J. Schnitzer. Emergent reliability in sensory cortical coding and inter-area communication. Nature, 605(7911):713–721, 2022.

T. Feng, D. Silva, and D. J. Foster. Dissociation between the experience-dependent development of hippocampal theta sequences and single-trial phase precession. Journal of Neuroscience, 35(12):4890–4902, 2015. ISSN 0270-6474. doi: 10.1523/JNEUROSCI.2614-14.2015. URL https://www.jneurosci.org/content/35/12/4890

J. A. Gallego, M. G. Perich, R. H. Chowdhury, S. A. Solla, and L. E. Miller. Long-term stability of cortical population dynamics underlying consistent behavior. Nature neuroscience, 23(2):260–270, 2020.

R. A. Harshman et al. Foundations of the parafac procedure: Models and conditions for an” explanatory” multimodal factor analysis. 1970.

C. D. Harvey, P. Coen, and D. W. Tank. Choice-specific sequences in parietal cortex during a virtual-navigation decision task. Nature, 484(7392):62–68, 2012. ISSN 1476-4687. doi: 10.1038/nature10918. URL https://doi.org/10.1038/nature10918

J. A. Hennig, E. R. Oby, D. M. Losey, A. P. Batista, M. Y. Byron, and S. M. Chase. How learning unfolds in the brain: toward an optimization view. Neuron, 109(23):3720–3735, 2021.

I. B. L. Ibl, V. Aguillon-Rodriguez, D. Angelaki, H. Bayer, N. Bonacchi, M. Carandini, F. Cazettes, G. Chapuis, A. K. Churchland, Y. Dan, E. Dewitt, M. Faulkner, H. Forrest, L. Haetzel, M. Häusser, S. B. Hofer, F. Hu, A. Khanal, C. Krasniak, I. Laranjeira, Z. F. Mainen, G. Meijer, N. J. Miska, T. D. Mrsic-Flogel, M. Murakami, J.-P. Noel, A. Pan-Vazquez, C. Rossant, J. Sanders, K. Socha, R. Terry, A. E. Urai, H. Vergara, M. Wells, C. J. Wilson, I. B. Witten, L. E. Wool, and A. M. Zador. Standardized and reproducible measurement of decision-making in mice. eLife, 10:e63711, may 2021. ISSN 2050-084X. doi: 10.7554/eLife.63711.

I. B. L. Ibl, K. Banga, J. Benson, N. Bonacchi, S. A. Bruijns, R. Campbell, G. A. Chapuis, A. K. Churchland, M. F. Davatolhagh, H. D. Lee, M. Faulkner, F. Hu, J. Hunterberg, A. Khanal, C. Krasniak, G. T. Meijer, N. J. Miska, Z. Mohammadi, J.-P. Noel, L. Paninski, A. Pan-Vazquez, N. Roth, M. Schartner, K. Socha, N. A. Steinmetz, M. Taheri, A. E. Urai, M. Wells, S. J. West, M. R. Whiteway, and O. Winter. Reproducibility of in-vivo electrophysiological measurements in mice. bioRxiv, 2022. doi: 10.1101/2022.05.09.491042. URL https://www.biorxiv.org/content/early/2022/05/09/2022.05.09.491042.1.

M. Jazayeri and S. Ostojic. Interpreting neural computations by examining intrinsic and embedding dimensionality of neural activity. Current opinion in neurobiology, 70:113–120, 2021.

D. P. Kingma and J. Ba. Adam: A method for stochastic optimization, 2014. URL https://arxiv.org/abs/1412.6980

S. A. Koay, A. S. Charles, S. Y. Thiberge, C. D. Brody, and D. W. Tank. Sequential and efficient neural-population coding of complex task information. Neuron, 110(2):328–349.e11, 2022. ISSN 0896-6273. doi: https://doi.org/10.1016/j.neuron.2021.10.020. URL https://www.sciencedirect.com/science/article/pii/S0896627321008357

D. Kobak, W. Brendel, C. Constantinidis, C. E. Feierstein, A. Kepecs, Z. F. Mainen, X.-L. Qi, R. Romo, N. Uchida, and C. K. Machens. Demixed principal component analysis of neural population data. Elife, 5:e10989, 2016.

T. G. Kolda and B. W. Bader. Tensor decompositions and applications. SIAM review, 51(3):455–500, 2009.

K. V. Kuchibhotla, T. Hindmarsh Sten, E. S. Papadoyannis, S. Elnozahy, K. A. Fogelson, R. Kumar, Y. Boubenec, P. C. Holland, S. Ostojic, and R. C. Froemke. Dissociating task acquisition from expression during learning reveals latent knowledge. Nature communications, 10(1):1–13, 2019.

K. C. Lakshmanan, P. T. Sadtler, E. C. Tyler-Kabara, A. P. Batista, and B. M. Yu. Extracting low-dimensional latent structure from time series in the presence of delays. Neural Comput., 27(9):1825–1856, sep 2015. ISSN 0899-7667. doi: 10.1162/NECO\-a\_00759.

F. Lanore, N. A. Cayco-Gajic, H. Gurnani, D. Coyle, and R. A. Silver. Cerebellar granule cell axons support high-dimensional representations. Nature Neuroscience, 24(8):1142–1150, 2021.

G. W. Lindsay, T. D. Mrsic-Flogel, and M. Sahani. Bio-inspired neural networks implement different recurrent visual processing strategies than task-trained ones do. bioRxiv, 2022.

E. L. Mackevicius, A. H. Bahle, A. H. Williams, S. Gu, N. I. Denisenko, M. S. Goldman, and M. S. Fee. Unsupervised discovery of temporal sequences in high-dimensional datasets, with applications to neuroscience. Elife, 8:e38471, 2019.

L. McInnes, J. Healy, and J. Melville. Umap: Uniform manifold approximation and projection for dimension reduction. arXiv preprint arXiv:1802.03426, 2018.

T. S. Okubo, E. L. Mackevicius, H. L. Payne, G. F. Lynch, and M. S. Fee. Growth and splitting of neural sequences in songbird vocal development. Nature, 528(7582):352–357, 2015. ISSN 1476-4687. doi: 10.1038/nature15741. URL https://doi.org/10.1038/nature15741

A. Onken, J. K. Liu, P. C. R. Karunasekara, I. Delis, T. Gollisch, and S. Panzeri. Using matrix and tensor factorizations for the single-trial analysis of population spike trains. PLoS computational biology, 12(11): e1005189, 2016.

C. Pandarinath, D. J. O’Shea, J. Collins, R. Jozefowicz, S. D. Stavisky, J. C. Kao, E. M. Trautmann, M. T. Kaufman, S. I. Ryu, L. R. Hochberg, et al. Inferring single-trial neural population dynamics using sequential auto-encoders. Nature methods, 15(10):805–815, 2018.

S. Panzeri, M. Moroni, H. Safaai, and C. D. Harvey. The structures and functions of correlations in neural population codes. Nature Reviews Neuroscience, pages 1–17, 2022.

N. F. Parker, A. Baidya, J. Cox, L. M. Haetzel, A. Zhukovskaya, M. Murugan, B. Engelhard, M. S. Goldman, and I. B. Witten. Choice-selective sequences dominate in cortical relative to thalamic inputs to nac to support reinforcement learning. Cell Reports, 39(7):110756, 2022. ISSN 2211-1247. doi: https://doi.org/10.1016/j.celrep.2022.110756. URL https://www.sciencedirect.com/science/article/pii/S2211124722005204.

E. Pastalkova, V. Itskov, A. Amarasingham, and G. Buzsáki. Internally generated cell assembly sequences in the rat hippocampus. Science, 321(5894):1322–1327, 2008. doi: 10.1126/science.1159775. URL https://www.science.org/doi/abs/10.1126/science.1159775.

A. Paszke, S. Gross, F. Massa, A. Lerer, J. Bradbury, G. Chanan, T. Killeen, Z. Lin, N. Gimelshein, L. Antiga, A. Desmaison, A. Kopf, E. Yang, Z. DeVito, M. Raison, A. Tejani, S. Chilamkurthy, B. Steiner, L. Fang, J. Bai, and S. Chintala. Pytorch: An imperative style, high-performance deep learning library. In Advances in Neural Information Processing Systems 32, pages 8024–8035. Curran Associates, Inc., 2019.

F. Pedregosa, G. Varoquaux, A. Gramfort, V. Michel, B. Thirion, O. Grisel, M. Blondel, P. Prettenhofer, R. Weiss, V. Dubourg, J. Vanderplas, A. Passos, D. Cournapeau, M. Brucher, M. Perrot, and E. Duchesnay. Scikit-learn: Machine learning in Python. Journal of Machine Learning Research, 12:2825–2830, 2011.

F. Pei, J. Ye, D. M. Zoltowski, A. Wu, R. H. Chowdhury, H. Sohn, J. E. O’Doherty, K. V. Shenoy, M. T. Kaufman, M. Churchland, M. Jazayeri, L. E. Miller, J. Pillow, I. M. Park, E. L. Dyer, and C. Pandarinath. Neural latents benchmark ‘21: Evaluating latent variable models of neural population activity. In Advances in Neural Information Processing Systems (NeurIPS), Track on Datasets and Benchmarks, 2021. URL https://arxiv.org/abs/2109.04463.

A. J. Peters, S. X. Chen, and T. Komiyama. Emergence of reproducible spatiotemporal activity during motor learning. Nature, 510(7504):263–267, 2014. ISSN 1476-4687. doi: 10.1038/nature13235. URL https://doi.org/10.1038/nature13235.

J. Poort, A. G. Khan, M. Pachitariu, A. Nemri, I. Orsolic, J. Krupic, M. Bauza, M. Sahani, G. B. Keller, T. D. Mrsic-Flogel, et al. Learning enhances sensory and multiple non-sensory representations in primary visual cortex. Neuron, 86(6):1478–1490, 2015.

A. Renart and C. K. Machens. Variability in neural activity and behavior. Current opinion in neurobiology, 25:211–220, 2014.

M. E. Rule, T. O’Leary, and C. D. Harvey. Causes and consequences of representational drift. Current Opinion in Neurobiology, 58:141–147, 2019.

E. Rybakken, N. Baas, and B. Dunn. Decoding of neural data using cohomological feature extraction. Neural computation, 31(1):68–93, 2019.

O. G. Sani, H. Abbaspourazad, Y. T. Wong, B. Pesaran, and M. M. Shanechi. Modeling behaviorally relevant neural dynamics enabled by preferential subspace identification. Nature Neuroscience, 24(1): 140–149, 2021a.

O. G. Sani, B. Pesaran, and M. M. Shanechi. Where is all the nonlinearity: flexible nonlinear modeling of behaviorally relevant neural dynamics using recurrent neural networks. bioRxiv, 2021b. doi: 10.1101/2021.09.03.458628. URL https://www.biorxiv.org/content/early/2021/09/06/2021.09.03.458628

M. Schimel, T.-C. Kao, K. T. Jensen, and G. Hennequin. ilqr-vae: control-based learning of input-driven dynamics with applications to neural data. bioRxiv, pages 2021–10, 2022.

C. E. Schoonover, S. N. Ohashi, R. Axel, and A. J. Fink. Representational drift in primary olfactory cortex. Nature, 594(7864):541–546, 2021.

W. Schultz. Predictive reward signal of dopamine neurons. Journal of neurophysiology, 80(1):1–27, 1998.

J. S. Seely, M. T. Kaufman, S. I. Ryu, K. V. Shenoy, J. P. Cunningham, and M. M. Churchland. Tensor analysis reveals distinct population structure that parallels the different computational roles of areas m1 and v1. PLoS computational biology, 12(11):e1005164, 2016.

K. V. Shenoy, M. Sahani, M. M. Churchland, et al. Cortical control of arm movements: a dynamical systems perspective. Annu Rev Neurosci, 36(1):337–359, 2013.

M. A.-Y. Smith, K. S. Honegger, G. Turner, and B. de Bivort. Idiosyncratic learning performance in flies. Biology Letters, 18(2):20210424, 2022.

C. Stringer, M. Pachitariu, N. Steinmetz, M. Carandini, and K. D. Harris. High-dimensional geometry of population responses in visual cortex. Nature, 571(7765):361–365, 2019.

T. Tao and W. Sawin. Notes on the “slice rank” of tensors, 08 2016. URL https://terrytao.wordpress.com/2016/08/24/notes-on-the-slice-rank-of-tensors/.

M. Vinck, R. Batista-Brito, U. Knoblich, and J. A. Cardin. Arousal and locomotion make distinct contributions to cortical activity patterns and visual encoding. Neuron, 86(3):740–754, 2015.

M. J. Wagner, T. H. Kim, J. Kadmon, N. D. Nguyen, S. Ganguli, M. J. Schnitzer, and L. Luo. Shared cortex-cerebellum dynamics in the execution and learning of a motor task. Cell, 177(3):669–682, 2019.

A. H. Williams, T. H. Kim, F. Wang, S. Vyas, S. I. Ryu, K. V. Shenoy, M. Schnitzer, T. G. Kolda, and S. Ganguli. Unsupervised discovery of demixed, low-dimensional neural dynamics across multiple timescales through tensor component analysis. Neuron, 98(6):1099–1115, 2018.

A. H. Williams, E. Kunz, S. Kornblith, and S. Linderman. Generalized shape metrics on neural representations. In M. Ranzato, A. Beygelzimer, Y. Dauphin, P. Liang, and J. W. Vaughan, editors, Advances in Neural Information Processing Systems, volume 34, pages 4738–4750. Curran Associates, Inc., 2021. URL https://proceedings.neurips.cc/paper/2021/file/252a3dbaeb32e7690242ad3b556e626b-Paper.pdf.

S. Zhou, S. C. Masmanidis, and D. V. Buonomano. Neural sequences as an optimal dynamical regime for the readout of time. Neuron, 108(4):651–658.e5, 2020. ISSN 0896-6273. doi: https://doi.org/10.1016/j.neuron.2020.08.020.URL https://www.sciencedirect.com/science/article/pii/S0896627320306516.

